# MARVEL: An integrated alternative splicing analysis platform for single-cell RNA sequencing data

**DOI:** 10.1101/2022.08.25.505258

**Authors:** Wei Xiong Wen, Adam J Mead, Supat Thongjuea

## Abstract

Alternative splicing is an important source of heterogeneity in gene expression between individual cells but remains an understudied area due to the paucity of computational tools to analyze splicing dynamics at single-cell resolution. Here, we present MARVEL, a comprehensive R package for single-cell splicing analysis applicable to RNA sequencing generated from the plate- and droplet-based methods. We performed extensive benchmarking of MARVEL against available tools and demonstrated its utility by analyzing iPSC differentiation into endoderm cells and cardiomyocytes. MARVEL enables systematic and integrated splicing and gene expression analysis of single cells to characterize the splicing landscape and reveal biological insights.

## INTRODUCTION

Single-cell RNA sequencing (scRNA-seq) is a powerful tool for studying transcriptional heterogeneity in normal tissues (1–5) and pathological conditions (6–11). The vast majority of scRNA-seq analyses focus on gene-level expression, however, alternative splicing represents an important additional layer of transcriptional complexity underlying gene expression (12). Alternative splicing has not been widely investigated at single-cell resolution and thus remains an untapped source of knowledge in both health and disease states (13). This is potentially due to the lack of available computational tools to interrogate alternative splicing for scRNA-seq data. Although existing analysis pipelines such as Seurat (14), Monocle (15), and Scanpy (16) enabled integrative analysis workflows for single-cell gene expression, they do not support comprehensive analyses to combine gene-level and alternative splicing information.

Recently, analysis tools, such as BRIE (versions 1 and 2) (17, 18) and Expedition (19), were developed to analyze alternative splicing in scRNA-seq datasets generated from the plate-based platforms e.g., Smart-seq2 (20). BRIE uses a Bayesian approach to learn informative sequence features for percent spliced-in (PSI) estimation, leading to the improvement of PSI estimation for splicing events that have low-to-no coverage in scRNA-seq data (17, 18). Expedition introduces the concept of “modalities” to stratify PSI distributions into discrete categories (19). However, a number of functionalities required to comprehensively characterize alternative splicing dynamics at the single-cell level are not yet available. For instance, current analysis tools focus on PSI quantification for skipped-exons (SE) and mutually exclusive exons (MXE) splicing events (17, 19) but did not include retained-introns (RI), alternative 5’ and 3’ splice sites (A5SS and A3SS), and alternative first and last exons (AFE and ALE). While SE are the major splicing event type (21), other types of splicing events are also important sources of gene expression heterogeneity and have been shown to contribute to the cellular phenotype. For example, RI are a source of neoantigens in melanoma (22), whereas A5SS, A3SS, AFE, and ALE are often dysregulated in myelodysplastic syndrome (MDS) and acute myeloid leukemia (AML) patients carrying mutations in genes encoding for splicing factors (21,23,24).

Modality classification enables the changes in splicing patterns across different cell populations (19). Biases from PCR amplification and library preparation prevalent in scRNA-seq have been shown to lead to a high proportion of false positives, in particular for the bimodal classification (25). Therefore, modality assignment should incorporate these technical biases to enable better classification of splicing patterns.

Taken together, current computational tools may not comprehensively facilitate the characterization of alternative splicing dynamics at single-cell resolution. Moreover, existing analysis workflows do not integrate gene expression and alternative splicing information into a single framework. Here, we introduce MARVEL, an R package for integrative single-cell alternative splicing and gene expression analysis. We benchmarked MARVEL against existing computational tools for single-cell alternative splicing analysis and demonstrated its utility by analyzing publicly available datasets generated from the plate- and droplet-based library preparation methods derived from induced pluripotent stem cells (iPSCs) differentiated into endoderm and cardiomyocytes, respectively (26, 27).

## MATERIALS AND METHODS

### Processing of publicly available datasets

#### Plate-based scRNA-seq datasets

To assess and validate the performance of MARVEL on scRNA-seq data generated from plate-based library preparation protocols, we retrieved four datasets from previous studies (15,19,26,28). Raw sequencing reads (FASTQ) were downloaded from the Sequence Reads Archive (SRA). Adapters and 3’ bases with Phred quality scores <20 were trimmed using Trim Galore 0.6.5 (29). Trimmed reads were mapped to the GRCh38 reference genome using STAR 2.6.1d in 2-pass mode (30). STAR was also used to detect and quantify splice junction counts, while RSEM v1.2.31 was used to quantify gene expression in transcripts per million (TPM). Binary Alignment Map (BAM) file statistics including total mapped reads and mitochondrial reads were computed using Samtools 1.9 (31).

The first dataset consisted of human-induced pluripotent stem cells (iPSCs), neural progenitor cells (NPCs), and motor neurons (MNs) (19). Cells with >100,000 mapped reads, >70% alignment rate, and <15% mitochondrial reads were retained for data with paired-end reads. For the dataset with single-end reads, cells with >5,000,000 mapped reads, >90% alignment rate, and <10% mitochondrial reads were retained (Supplementary Figures 1A-F). Single cells that were annotated as outliers by the original study were excluded. In total, 62 iPSCs, 68 NPCs, and 60 MN cells were included for analysis. In addition, 2 iPSC, 3 NPC, and 3 MN matched-bulk samples were included for analysis.

The second dataset consisted of human myoblasts cultured and sequenced at 0-, 24-, 48-, and 72-hour (15). Cells with >100,000 mapped reads, >75% alignment rate, and <20% mitochondrial reads were retained (Supplementary Figures 1G-I). Single cells that were annotated as control wells by the original study were excluded. In total, 82, 85, 88, and 72 myoblasts at 0-, 24-, 48-, and 72-hour time points, respectively, were included for analysis. In addition, 3 matched-bulk samples for each time point were included for analysis.

The third dataset consisted of iPSC and endoderm cells (26). Cells with >100,000 mapped reads, >75% alignment rate, and <20% mitochondrial reads were retained (Supplementary Figures 1J-L). Five cells that were annotated as the unknown cell type by the original study were excluded. In total, 83 iPSC and 53 endoderm cells were included for analysis.

The fourth dataset consisted of single cells derived from the spinal cord of mice induced with experimental autoimmune encephalomyelitis (EAE) and control mice (28). Cells that passed sequencing QC were defined as having read alignment > 50%, > 40,000 mapped reads, and mitochondrial reads < 55% (Supplementary Figures 1M-O). Eight cells annotated as doublets by the original publication were removed. In total, 1,078 EAE and 978 control mice cells were included for analysis.

All datasets were used for benchmarking MARVEL. The third dataset consisting of iPSC and endoderm cells was used to demonstrate the analyses provided by MARVEL.

#### 10x Genomics dataset

To demonstrate the utility of MARVEL for single-cell alternative splicing on a dataset from a droplet-based platform, we retrieved scRNA-seq data from iPSC and iPSC-derived cardiomyocytes on days 2, 4, and 10 generated using 10x Genomics Chromium Single Cell 3’ Reagent Kit (version 2) (27). Raw sequencing reads (FASTQ) were downloaded from the Sequence Reads Archive (SRA) and were aligned to the GRCh38 reference genome using Cell Ranger v2.1.1. The resulting BAM files for each sample were used as inputs for STARsolo (available in STAR v2.7.8a) to generate the gene expression count matrices (32). SingCellaR (1, 33) was subsequently used to identify and retain good-quality cells based on per-cell UMI counts and the number of detected genes (Supplementary Figures 2A-D) (1). Additionally, only cells with <15% mitochondrial counts and genes expressed in at least 10 cells were retained. Good-quality cells from iPSC and day-10 cardiomyocytes were subsequently integrated and the t-Distributed Stochastic Neighbor Embedding (tSNE) coordinates were generated using SingCellaR.

### Isoform detection

#### Plate-based scRNA-seq datasets

We analyzed isoform usage at the exon level for scRNA-seq data. For each cell type, the bulk samples were used to create a cell-type-specific gene transfer file (GTF) using StringTie2 (34). When the bulk samples were not available, pseudo-bulk samples were generated by merging the single-cell BAM files. The GENCODE GTF v31 file was used as a guide to generate the cell-type-specific GTF files (35). The GTF represents the transcriptome assembly and hence the catalog for all genes, transcripts, and exons detected for a particular cell type. The cell-type-specific GTF files were then merged to obtain the final GTF file for isoform detection in BRIE and MARVEL. Expedition performed *de novo* detection of splice junctions, exons, and alternative splicing events directly on the single-cell and bulk BAM files.

MARVEL takes SE, MXE, RI, A5SS, and A3SS splicing events detected by rMATS 4.1.0 (36). rMATS identifies these alternative splicing events using the aforementioned merged GTF file as previously described (25). Splice junction counts were generated using STAR 2.6.1d in a two-pass mode (30). The gene expression matrix, splicing junction count matrix, detected exon-level alternative splicing events, and GENCODE GTF v31 file, were used as inputs for MARVEL. MARVEL created an R object from these inputs using the *CreateMarvelObject* function for downstream data processing and analyses. After creating the MARVEL object, additional splicing event types, AFE and ALE, were detected by MARVEL from the GENCODE GTF v31 provided by using the *DetectEvents* function.

#### 10x Genomics dataset

We analyzed isoform usage at the splice junction level for scRNA-seq data generated from a droplet-based platform. Splice junction counts were generated using STAR v2.7.8a (STARsolo). The filtered gene count matrix and tSNE coordinates from SingleCellaR, raw splice junction count from STARsolo, and reference gene transfer file (GTF) were used as inputs for MARVEL. MARVEL created an R object from inputs using the *CreateMarvelObject.10x* function for downstream data processing and analyses.

### Isoform validation and quantification

#### Plate-based scRNA-seq datasets

The percent spliced-in (PSI) values were used to measure the degree of alternative exon inclusion for scRNA-seq data generated from the plate-based protocol. The *briekit-event* function in BRIE was used to detect alternative splicing events from the GTF provided (see “Isoform detection” section). Next, the *briekit-event-filter* function was used together with the options *[--add_chrom chrX --as_exon_min 10 -- as_exon_max 100000000 --as_exon_tss 10 --as_exon_tts 10 --no_splice_site*] to filter for high-quality alternative splicing events. The *briekit-factor* function was then used to calculate the set of sequence features to use to infer PSI values for each alternative splicing event. Only skipped-exon (SE) splicing event was analyzed by BRIE here. The PSI values of these detected splicing events were subsequently quantified in three different modes using the *brie-quant* function with the options *--interceptMode None*, *--interceptMode cell*, and *--interceptMode gene*. The first mode (mode 0) uses a prior distribution centered at 0.5 to impute PSI values for alternative splicing events. The second mode (mode 1) combines a prior distribution centered at 0.5 with an informative prior inferred from genomic sequence-based features to impute PSI values. The third mode (mode 2) uses a prior distribution centered on the mean PSI values across the cell population to impute PSI values.

For Expedition analysis, we performed *de novo* detection of alternative splicing events using the *outrigger index* function and computed the PSI values for each alternative splicing event using the *outrigger psi* function. Each PSI value represents the fraction of splice junction reads supporting the alternative exons over the total splice junction reads supporting or skipping the alternative exons. For each cell, alternative splicing events supported by <10 splice junction reads were annotated as missing values. The types of alternative splicing events analyzed by Expedition included SE and MXE.

The types of exon-level alternative splicing events analyzed by MARVEL included seven main exon-level alternative splicing events comprising SE, MXE, RI, A5SS, A3SS, AFE, and ALE. To ensure high-quality alternative splicing events for downstream analyses, both alternative and constitutive exons of these alternative splicing events needed to be supported by the splice junction reads. Only alternative splicing events whose exons were supported by splice junction reads were retained (Supplementary Figure 3A). Furthermore, for the RI event, introns that overlap with any alternative or constitutive exons were filtered away (37). This resulted in high-quality introns and was termed “independent” intron because they do not overlap with any annotated exons.

Similar to Expedition, MARVEL uses a splice junction-based approach to compute PSI values. For SE, MXE, A5SS, A3SS, AFE, and ALE alternative splicing events, PSI values are computed as a fraction of splice junction reads supporting the alternative exons over the total splice junction reads supporting or skipping the alternative exons (Supplementary Figures 3B-G).

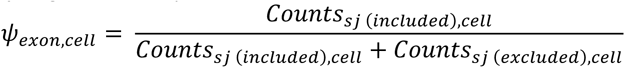

For the RI event, the PSI values of the intron are computed as the total intron read counts normalized by the intronic length and then divided by the sum of total intron read counts normalized by the intronic length and total splice junction counts skipping the introns (Supplementary Figure 3H).

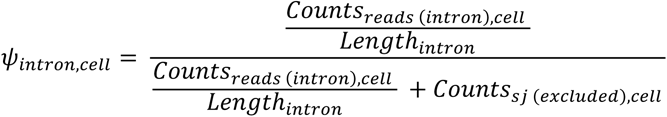

The *ComputePSI* function was used to validate the alternative splicing events and calculate the corresponding PSI values. SE, MXE, A5SS, A3SS, AFE, and ALE alternative splicing events supported by <10 of splice junction reads supporting or skipping the alternative exons in a given cell were annotated as missing values. RI alternative splicing events supported by <10 of length-normalized introns read counts or <10 of splice junction skipping introns in a given cell were annotated as missing values.

#### 10x Genomics dataset

To ensure the inclusion of high-quality splicing junctions for downstream analyses, the exons of each splice junction were cross-checked with the GTF file and were categorized as annotated, multi-mapped, and unannotated. An annotated exon is an exon that maps to a single gene. A multi-mapped exon is an annotated exon that maps to multiple genes. An unannotated exon is an exon with no matching record in the GTF file. Only splice junctions consisting of both annotated exons were retained for downstream analyses. The *AnnotateSJ.10x* function was used to annotate each splice junction, while the *FilterSJ.10x* function was used to filter splicing junctions consisting of unannotated and multi-mapped exons.

Splice junction usage was used to measure the degree of splice junction inclusion. For a given cell type, the splice junction usage was computed as the fraction of the sum of splice junction counts across all cells over the sum of gene counts for the corresponding splice junction across all cells (38).

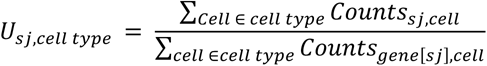

The *ComputeSJusage.10x* function was used to compute the cell type-specific usage of validated splice junctions.

### Benchmarking processing time for isoform quantification

To compare processing time between BRIE and MARVEL, we measured the time taken to compute the PSI values for the same set of 1,000 SE splicing events. To compare processing time between Expedition and MARVEL, we measured the time taken to compute the PSI values for the same set of 500 SE and 500 MXE splicing events. We additionally measured the time taken to compute the PSI values for 1,000 splicing events per each of RI, A5SS, A3SS, AFE, and ALE. We measured the processing time using the Slurm Workload Manager (v20.02.0) on CentOS Linux 7 (Core).

### Sequence conservation analysis

The sequence conservation scores for the 5’ and 3’ constitutive exons, and alternative exons were computed using the *phastCons100way.UCSC.hg38* R package (39). The Pearson correlation between alternative exon conservation scores and mean PSI values for each cell line was analyzed (26).

### Modality assignment

Song *et al.* proposed that the PSI values for a given alternative splicing event can be categorized into five modalities comprising included, excluded, bimodal, middle, and multimodal (19). Here, MARVEL models each alternative splicing event as a beta distribution and estimates the alpha and beta parameters using the maximum likelihood approach. Based on the parameters’ values, each alternative splicing event was categorized sequentially into their respective modality as follows (see also Supplementary Figures 4A-E):

1. Bimodal (PSI ≈ 0, 100): α < 0.5 or β < 0.5

2. Included (PSI ≈ 100): α > β

3. Excluded (PSI ≈ 50): β < α

4. Middle (PSI ≈ 50): α > 1 & β > 1 & α = β

5. Multimodal (uniform distribution): α = β = 1

MARVEL further expands the current repertoire of modalities by stratifying the included and excluded modalities into primary and dispersed. In included and excluded primary modalities, the PSI values cluster tightly around 100 and 0. In included and excluded dispersed modalities, PSI values cluster towards 100 and 0 with the addition of some values that trended towards opposite ends. Therefore, the dispersed modality has a higher variance among PSI values than the primary modality. Here, we applied a heuristic threshold of variance at 0.001 to categorize the included and excluded modalities into primary (<0.001) and dispersed (>=0.001; Supplementary Figures 4B-C).

Modality assignment for alternative splicing events was performed using the *AssignModality* function. In this study, alternative splicing events supported by at least 10 reads in at least 25 cells were included for modality assignment by Expedition and MARVEL.

### Bimodality adjustment

A significant proportion of bimodal splicing patterns detected were previously reported to be artifacts of single-cell RNA-sequencing (25). To distinguish between true and false (spurious) bimodal splicing patterns, we generated a set of false and true positive bimodal splicing patterns from a previous study with experimental validation of alternative splicing events detected from RNA-sequencing using quantitative polymerase chain reaction (qPCR) and small molecular fluorescent *in situ* hybridization (smFISH) (19). We further expanded our search for additional true bimodal splicing patterns in two additional studies (15, 26) whereby bimodal splicing patterns constituted of single cells in which the corresponding gene had 10 or more mRNA molecules as previously described (25). The mRNA count for each gene was computed using the *monocle* R package (15). In total, 45 true and 7 false bimodal splicing patterns were included for analysis. We assessed the ability of three features to distinguish true from false bimodal patterns: (1) fold difference between the proportion of cells with PSI > 75 and PSI < 25 (and vice versa), (2) difference between the proportion of cells with PSI > 75 and PSI < 25 (and vice versa), and (3) average PSI value. Heuristic thresholds of 75 and 25 were chosen because they distinguished true from false bimodal patterns (Figures 2F-H). The argument *bimodal.adjust=TRUE* can be used in the *AssignModality* function to detect true and false and subsequently reassign false bimodality into either included or excluded modality.

We then assessed the ability of Expedition and MARVEL to distinguish bimodality and non-bimodality. From the datasets (15,19,26), we generated a set of alternative splicing events consisting of 45 bimodal and 17,259 non-bimodal (included, excluded, middle, and multimodal) modalities as previously described in (25). We cross-tabulated the bimodal and non-bimodal assignment of Expedition and MARVEL against this set of ground truths to create a confusion matrix. This allowed us to compute and compare several evaluation metrics consisting of sensitivity, specificity, negative predictive value, and precision for Expedition and MARVEL.

### Differential splicing analysis

#### Plate-based scRNA-seq datasets

To detect differences in splicing patterns between groups of single cells, we need to take into account both mean and variance. For example, it would not be possible to distinguish between bimodal, middle, and multimodal splicing patterns based on mean alone. To this end, MARVEL implements three nonparametric statistical tests for assessing the differences in splicing patterns between groups of single cells. These are the Kolmogorov-Smirnov, Anderson-Darling (AD) (40) and D Test Statistic (DTS) approaches (41). These tests take into account the overall PSI distribution and assess the differences in PSI distribution for each alternative splicing event between groups of single cells. MARVEL combines differential splicing analysis using AD and DTS, followed by the outlier removal. MARVEL also includes Wilcoxon rank-sum test, t-test, and permutation test for differential alternative splicing analysis, used for comparing PSI values in bulk samples. Differential splicing analysis can be performed using the *CompareValues* function and subsequently visualized using the *PlotDEValue* function. In this study, alternative splicing events supported by at least 10 reads in least at 25 cells were included for differential splicing analysis. Alternative splicing events with FDR < 0.10 were considered to be differentially spliced.

For BRIE, differential splicing analysis was performed using the *brie-quant* function with default parameters. Alternative splicing events with evidence of lower bound (ELBO) gain > 4 were considered to be differentially spliced as previously described (18).

#### 10x Genomics dataset

Due to the sparsity of genes and splice junctions detected in scRNA-seq data generated from a droplet-based method, we performed differentially splicing analysis at the cell type level (pseudo-bulk) as opposed to at the single-cell level (38, 42). MARVEL utilizes a permutation approach for assessing differentially spliced junctions between two cell populations (43). For a given splice junction, the cell-type labels of single cells in the two cell populations are shuffled (permutated). PSI values per cell population are computed. Differences in the PSI values between populations are noted (ΔPSI_permutated_). The differential process is iterated 100 times, and these differences in the PSI values will form the null distribution. Then, the observed differences in the PSI values between the cell populations (ΔPSI_observed_) are compared against the null distribution to obtain *P*-values. Differentially spliced junctions were defined as mean log2-transformed normalized gene expression > 1.0, |Δ PSI| > 5, and *P*-value < 0.05. Differential splicing analysis can be performed using the *CompareValues.PSI.10x* function and subsequently visualized on the PCA/tSNE/UMAP plot using the *PlotDEValues.PCA.10x* function.

To assess the ability of MARVEL to identify biologically relevant genes that are differentially spliced, we performed differential splicing analysis on a dataset generated from iPSC and iPSC-derived cardiomyocytes on day 10 (27).

### Differential gene expression analysis

For plate-based scRNA-seq datasets, differential gene expression analysis was performed using the *CompareValues.Exp* function and subsequently visualized using the *PlotDEValue.Exp* function in MARVEL. Wilcoxon rank-sum test was used to assess the differences in normalized and log2-transformed gene expression values between two cell populations. Genes with FDR < 0.10 and log2 fold change of > 0.5 or < -0.5 were considered to be differentially expressed.

For a droplet-based scRNA-seq dataset, differential gene expression analysis was performed using the *CompareValues.Exp.Global.10x* function and subsequently visualized using the *PlotDEValues.Exp.Global.10x* function in MARVEL. Wilcoxon rank-sum test was used to assess the differences in normalized and log2-transformed gene expression values between two cell populations. Genes with FDR < 0.10 and log2 fold change of > 1 or < -1 were considered to be differentially expressed.

### Gene ontology analysis

MARVEL implements the gene ontology analysis provided by the *clusterProfiler* R package (44, 45). Gene ontology analysis to detect enriched pathways among differentially spliced genes can be performed using the *BioPathways* and *BioPathways.10x* functions for scRNA-seq data from the plate- and droplet-based methods.

### Linear dimension reduction analysis

For plate-based analysis, principal component analysis (PCA) was used for linear dimension reduction analysis. Only genes and alternative splicing events expressed in at least 3 and 25 cells, respectively, were included for analysis. For alternative splicing events whose PSI values were NA, i.e., coverage <10 in a given cell, they were re-coded randomly with values ranging from 0-100 prior to dimension reduction analysis. PCA was performed and visualized by MARVEL using the *RunPCA* function.

### Nonsense-mediated decay (NMD) prediction

For a given alternative exon with >5 PSI difference between iPSCs and endoderm cells and FDR < 0.10, MARVEL will retrieve the gene identifier from which the alternative exon is related. All protein-coding isoforms from this gene that encode the alternative exon are retrieved. MARVEL inserts the alternative exon sequence into these isoforms and predicts the resulting amino acid sequences using the *translate* function implemented by the *Biostrings* R package. The position(s) of any stop codon and its relative position in base-pair to the final exon-exon junction is noted. Consequently, there are four categories of isoforms (1) alternative exons belonging to novel isoforms (no matching record in GTF), (2) non-protein-coding isoforms (isoforms with no open reading frame), (3) protein-coding isoforms with a premature terminal codon (PTC) introduced by the alternative exons, and (4) protein-coding isoforms whose open reading frame are not disrupted by the alternative exons.

Protein-coding isoforms are further stratified into isoforms that are subjected to nonsense-mediate decay (NMD) or not (21). For the former, PTC(s) are located within 50bp before the final exon-exon junction. For the latter, the isoforms either have PTC(s) located further than 50bp before the final exon-exon junction or the isoforms do not have any PTC(s) introduced by the alternative exons.

## RESULTS

### MARVEL provides a comprehensive alternative splicing analysis framework for scRNA-seq generated from plate-based and 10x Genomics platforms

The workflow requires pre-processed tab-delimited files to perform MARVEL analysis for scRNA-seq data generated from plate-based protocols. The input files include spliced junction count and normalized gene expression matrices, alternative splicing events, gene and sample metadata (Figure 1A). Spliced junction counts, normalized gene expression values, and alternative splicing events are generated by STAR aligner (32), RSEM (46), and rMATS (36), respectively. It is noteworthy that rMATS provides splicing events from alternative and constitutive exons that are mapped to the same gene, thus avoiding ambiguous multi-mapping exons, i.e., exons mapped to multiple genes. First, MARVEL identifies high-quality alternative splicing events, defined as events validated by at least 10 splice junctions reads (Figure 1B). The PSI values of these alternative splicing events will be subsequently computed for downstream analyses, using the total number of reads supporting the alternative exon divided by the total number of reads supporting the alternative exon and constitutive exons (Figure 1C). Only junction-spanning reads, but not reads aligning to exon bodies, are used for PSI quantification (19,47,48). MARVEL performs linear and non-linear dimensionality reduction based on PSI values using principal component analysis (PCA) (Figure 1D), t-distributed stochastic neighbor embedding (t-SNE) (49), and uniform manifold approximation and projection (UMAP) (50). To assign alternative splicing events into distinct modalities (splicing patterns based on PSI distributions comprising included, excluded, bimodal, and multimodal (19)), MARVEL models each PSI distribution as a beta distribution and corrects the bimodal distributions for PCR amplification and library preparation biases (Figure 1E). In addition to the original modalities prescribed previously (19), MARVEL further stratifies the included and excluded modalities into primary and dispersed, thus providing a finer classification of PSI distribution. MARVEL also performs differential splicing and gene expression analysis to reveal the changes in alternative splicing relative to gene expression in addition to changes in modality across different cell populations (Figure 1F). In addition, MARVEL provides a function for gene ontology analysis of differentially spliced genes to reveal gene sets that are coordinatedly spliced. Furthermore, to explore the effect of alternative splicing on gene expression, MARVEL can be used to predict nonsense-mediated decay (NMD) of differentially spliced genes (Figure 1G).

**Figure 1.**
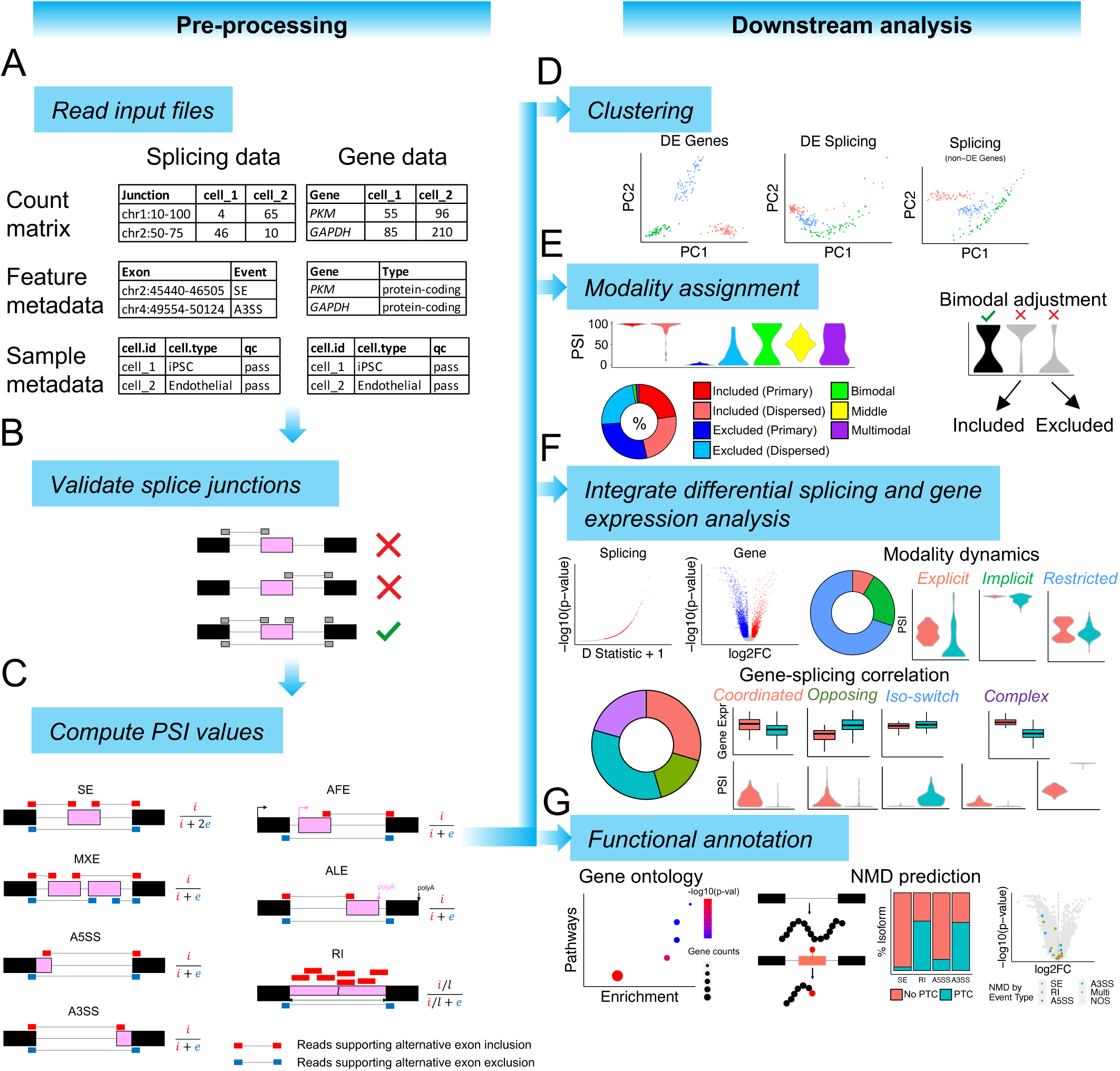
MARVEL workflow for single-cell alternative splicing analysis in RNA-sequencing dataset generated from plate-based methods. (**A-C**) Workflow for pre- processing of splicing and gene expression data by MARVEL. (**A**) Input files required by MARVEL include splice junction count and normalized gene expression matrix, alternative splicing events, and gene and sample metadata. (**B**) Only alternative splicing events supported by at least 10 splice junction reads are retained. (**C**) The PSI values of the confident alternative splicing events identified in (**B**) are computed for main exon-level alternative splicing event types. PSI values are calculated as the total number of reads supporting the alternative exons (pink) divided by the total number of reads supporting both alternative exons and constitutive exons (black). (**D-G**) Downstream analyses using computed PSI and gene expression values. (**D**) Dimension reduction analysis using differentially expressed genes (left), PSI values (middle), and PSI values of non-differentially expressed genes (right). (**E**) The assignment of PSI distributions into seven modalities (as indicated by colors) and the bimodal classification adjustment to reduce false bimodal classification. (**F**) Differential splicing and gene expression analysis and characterization of the alternative splicing in different modality changes or relative to gene expression changes across different cell populations. (**G**) Pathway enrichment analysis of differentially spliced genes and NMD prediction of alternative splicing events to understand the functional consequences of differential alternative splicing events. Genes subjected to NMD are visualized on the volcano plot generated from differential gene expression analysis. A3SS: alternative 3’ splice site; A5SS: alternative 5’ splice site; AFE: Alternative first exon; ALE: Alternative last exon; DE: Differentially expressed; FC: Fold change; iPSC: Induced pluripotent stem cell; MN: Motor neuron; MXE: mutually exclusive exons; NMD: Nonsense-mediated decay; NPC: Neural progenitor cell; PC: Principal component; PSI: percent spliced-in; PTC: Premature terminal codon; RI: retained-intron; SE: skipped-exon

Similar to plate-based alternative splicing analysis, MARVEL requires pre-processed tab-delimited files as input for droplet-based (e.g., 10x Genomics) scRNA-seq data analysis. MARVEL takes the sparse splice junction and gene count matrices, normalized gene expression matrix, and splice junction, gene and sample metadata (Supplementary Figure 5A). Splice junction and gene counts are generated by STARsolo (38), and normalized gene expression values are generated by SingCellaR (33). In this step, the cell-level information, such as pre-defined cell types, and the coordinates of dimension reduction embeddings, analyzed using other pipelines, can be incorporated. MARVEL identifies and removes splice junctions comprising multi-mapping exons (exons mapping to multiple genes) and retains only uniquely mapped exons (Supplementary Figure 5B). The splice junction usage of the remaining high-quality splice junctions is computed as the total splice junction counts divided by the total corresponding gene counts (Supplementary Figure 5C) (38). MARVEL allows users to explore the splice junction and gene expression distribution and select informative (expressed) genes and splice junctions for downstream analysis (Supplementary Figure 5D). MARVEL integrates differential splice junction usage and gene expression analysis and performs functional annotation of differentially spliced genes to identify gene sets that are coordinatedly spliced (Supplementary Figures 5E and F). We summarize analysis features provided by MARVEL compared to available alternative splicing computational tools described in Supplementary Table 1.

### MARVEL is benchmarked against established packages

#### Percent spliced-in estimation

To estimate PSI values from scRNA-Seq data, Bayesian regression prediction- and sequencing read-based approaches have been used (13,17–19,26). The former incorporates genomic features such as nucleotide context and cell-specific features such as cell-type, together with sequencing reads into the PSI value prediction, whereas the latter uses only sequencing reads, specifically splice junction reads, to compute PSI values.

The Bayesian approach based on genomic features for PSI estimation has been applied to SE splicing events only (17). Here, we assessed the predictive value of a genomic feature in inferring PSI values for other types of splicing events. We observed relatively low-level correlations between PSI values from MXE, RI, A5SS, A3SS, AFE, and ALE splicing events and the phastCons conservation scores compared to SE (Figures 2A-B). The phastCons score was identified as the most predictive feature for PSI value estimation using BRIE, as previously described (26). This would suggest reliable estimation of PSI values if the Bayesian regression based on genomic features was to be applied to SE, but other methods of PSI estimation, such as those based on sequencing reads alone, may be more suitable for non-SE splicing events. Therefore, MARVEL implements the splice junction read-based for PSI estimation of SE, MXE, RI, A5SS, A3SS, AFE, and ALE.

To assess the precision of estimated PSI values computed using a splice junction read-based approach, we evaluated the reproducibility of PSI quantification across homogenous cell populations in different cell types (51). Compared to all three modes of computing PSI values by BRIE, the median cell-to-cell correlation in PSI values for SE splicing event was higher for Expedition and MARVEL (Figure 2C). Expedition and MARVEL showed similar median cell-to-cell correlation in PSI values for SE and MXE splicing events. This is because Expedition and MARVEL both used splice junction counts to compute PSI values. In addition, MARVEL computes PSI values for RI, A5SS, A3SS, AFE, and ALE that are not provided by BRIE and Expedition. Furthermore, we assessed the median cell-to-bulk correlation for SE splicing events and observed a significantly higher correlation for MARVEL and Expedition compared to BRIE modes 0 and 1 (Figure 2D). There was no significant difference in median cell-to-bulk correlation in PSI values for SE and MXE splicing events between MARVEL and Expedition. The overall median cell-to-cell and cell-to-bulk correlation for SE, MXE, RI, A5SS, A3SS, AFE, and ALE splicing events computed by MARVEL were generally higher than 0.82 per category, suggesting robust PSI values generated using MARVEL.

**Figure 2.**
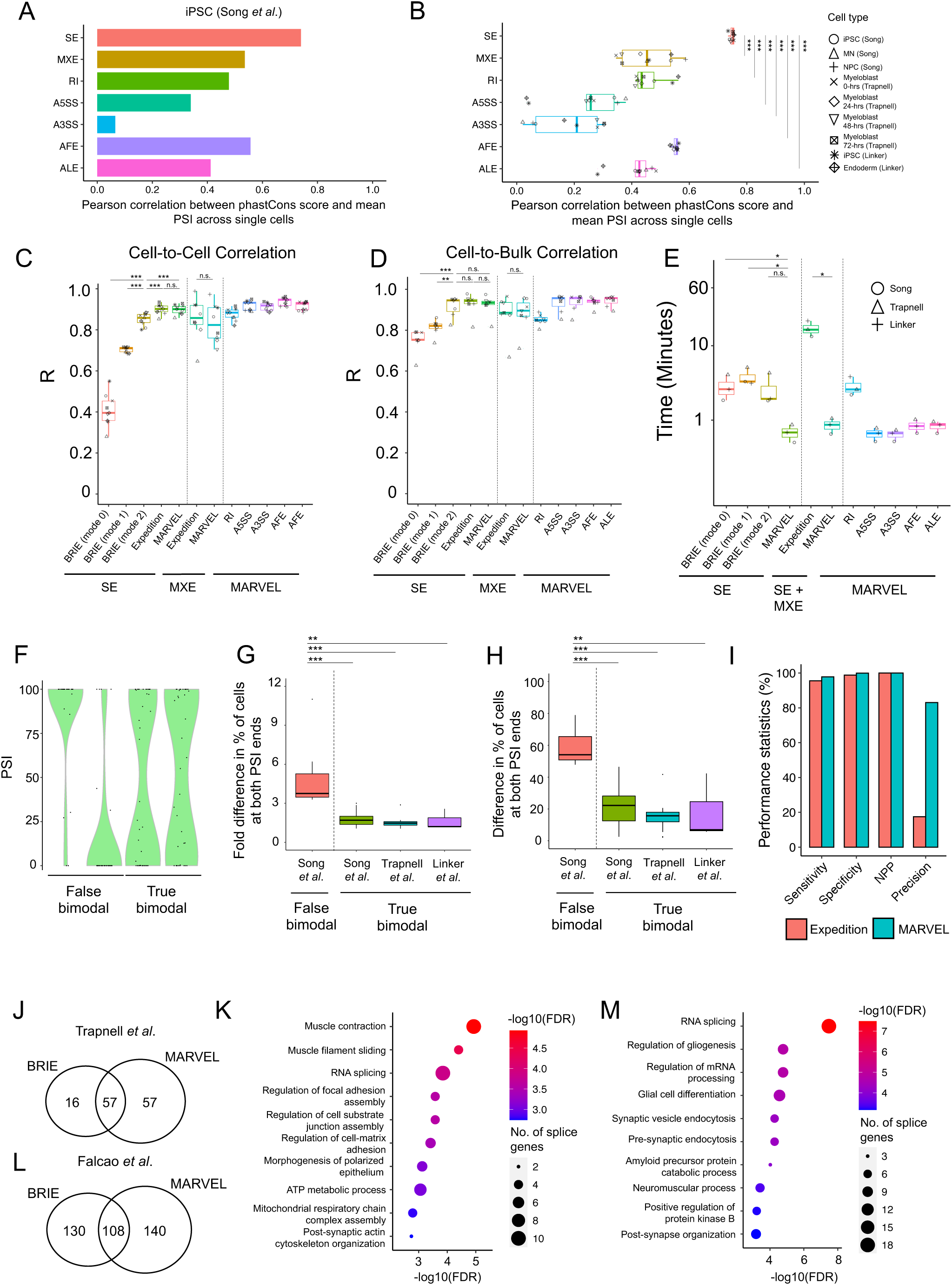
Benchmarking MARVEL against existing computational tools for single-cell alternative splicing analysis. **(A-B)** Pearson correlation between sequence conservation (phastCons) scores of alternative exons and their corresponding average PSI values across all single cells for each splicing event type in **(A)** iPSCs and **(B)** across nine cell lines. **(C)** Pearson correlation of PSI values between all possible single-cell pairs in nine cell lines compared across available splicing analysis tools. **(D)** Pearson correlation of PSI values between single cells and matched bulk sample in seven cell lines with both single cell and matched bulk sample available. **(E)** Comparison of processing time used to compute PSI values for 1,000 splicing events for each splicing event type in three datasets. **(F)** Representative false and true bimodal distributions from qPCR, smFISH, and mRNA count-based approach [15, 19, 26]. **(G-H)** Comparison of the **(G)** fold change (ratio) and **(H)** fold difference in the percentage of cells with PSI > 75 and PSI < 25 (and vice versa) between false and true bimodal distributions. **(I)** Evaluation metrics compared MARVEL versus Expedition for sensitivity, specificity, negative predictive value, and precision. The classification of false and true bimodal distributions predicted by MARVEL and Expedition was compared against the catalog of ground truths comprising 17,304 false and true bimodal distributions. **(J)** A Venn diagram showing the number of differentially spliced SE events detected between 72- vs. 0-hrs myoblast by BRIE and MARVEL. **(K)** MARVEL identified muscle-related pathways among differentially spliced genes between 72- vs. 0-hrs myoblast. **(L)** A Venn diagram showing the number of differentially spliced SE events detected between EAE and control mice using BRIE and MARVEL. **(M)** MARVEL identified neuron-related pathways among differentially spliced genes between EAE and control mice. hrs: hours; iPSC: induced pluripotent stem cell; MN: Motor neuron; NPC: Neural progenitor cell. *** FDR<0.01 ** FDR<0.05 * FDR<0.1.

We further compared the computational efficiency in processing time and Random-Access Memory (RAM) usage for computing the PSI values by BRIE, Expedition, and MARVEL. MARVEL required less time to compute the PSI values than BRIE and Expedition for the same dataset of SE and MXE splicing events (Figure 2E). Except for RI, the processing time across all datasets for A5SS, A3SS, AFE, and ALE was less than one minute. Lastly, the RAM usage was slightly higher for MARVEL than BRIE but lower than Expedition for computing the PSI values for the same dataset (Supplementary Figure 6A).

Taken together, MARVEL enables computationally efficient PSI quantification of all exon-level splicing event types and demonstrates reproducible PSI values across different cell populations that are comparable with existing single-cell splicing software.

#### Modality classification and correction

Modality classification concept was previously introduced to stratify the PSI distribution in a cell population into discrete categories, comprising excluded (PSI ∼ 0), included (PSI ∼ 100), middle (PSI ∼ 50), bimodal (PSI ∼ 0,100), and multimodal (uniform distribution) (19). However, the current modality assignment does not identify and correct for false classification caused by technical noise of scRNA-Seq experiments. Specifically for bimodal distributions, recent simulated and empirical data showed that a significant proportion of bimodality was spurious due to PCR amplification bias during the single-cell library preparation (25). Analyzing highly expressed alternative splicing events defined as genes with high mRNA count, and hence genes with low possibility of dropouts, e.g., genes with at least 10 molecules, has been shown to mitigate the false bimodal classification (25). However, this approach would preclude the majority of genes from downstream alternative splicing analysis. For example, we observed that >90% of genes were excluded when at least 10 molecules were required (Supplementary Figures 6B and C).

To retain alternative splicing events for analysis irrespective of gene abundance and at the same time mitigate false bimodal classification, we sought to identify distinguishable features between true and false bimodal distributions. We tabulated a set of true and false bimodal distributions encompassing alternative splicing events previously validated using qPCR or smFISH (19). Additionally, we tabulated a set of true bimodal distributions consisting of highly expressed alternative splicing events as previously described (25) (Figure 2F). We observed that the fold difference or difference in the proportion of cells at both ends of PSI distribution could delineate true from false bimodal distributions with thresholds of < 3 and < 50%, respectively (Figures 2G and H). Therefore, MARVEL incorporated these heuristic thresholds to identify true bimodality. Moreover, true bimodality revealed an average PSI (from both ends) of about 50, whereas the PSI values of false bimodality trended towards 100 or 0 (Supplementary Figure 6D). Therefore, MARVEL reclassifies the false bimodality into included or excluded modalities when the average PSI value is above or below 50.

Next, using our bimodal-adjusted modality approach, we compared Expedition’s and MARVEL’s ability to distinguish bimodal distributions from other modalities (e.g., included, excluded, middle, and multimodal). We tabulated a set of presumed true bimodal and non-bimodal distributions based on qPCR and smFISH validation and mRNA counts comprising 17,304 splicing events (25) (Supplementary Figure 6E). Expedition and MARVEL showed similar sensitivity, specificity, and negative predictive values in classifying bimodal and non-bimodal distributions (Figure 2I). However, Expedition showed a higher number of non-bimodality, leading to higher false-positive rates than MARVEL (*p* < 0.01; Fisher’s exact test; Supplementary Figure 6F). MARVEL, therefore, is more precise in classifying bimodality than Expedition.

Finally, we compared the percentage of splicing events classified as bimodal distribution by MARVEL and Expedition for highly expressed alternative splicing events and alternative splicing events that do not meet the criteria for high mRNA counts. Expedition classified a median of 7.8% of all splicing events as bimodal distribution compared to 1.4% by MARVEL, whereas only 0.2% of highly expressed splicing events were classified as bimodal distribution (Supplementary Figure 6G). However, MARVEL identified a bimodal distribution of the lowly expressed *PKM* gene, previously validated using smFISH (19). This gene would be missed if only highly expressed alternative splicing events were included for analysis (Supplementary Figures 6H and I). It is also noteworthy that almost four times more splicing events were eligible for modality assignment by MARVEL than when only highly expressed alternative splicing events were included for analysis.

Taken together, MARVEL leverages the concept of modality assignment introduced previously (19) while adjusting for technical biases from scRNA-seq library preparation (25) to enable robust classification of splicing patterns.

#### Differential splicing analysis

Statistical approaches for differential splicing analysis for scRNA-seq, such as BRIE (17), BRIE2 (18), and Expedition (19) were recently developed. However, BRIE requires a pairwise comparison between all possible pairs of cells that might use high computational resources and processing time to detect differential splicing events in a large number of cells (17). BRIE’s approach is suitable for detecting differential splicing events within a cell population but not across two different populations. Expedition detects differential splicing events based on modality change, excluding splicing events that show no modality change across cell populations (19). Detection of differential splicing events based on modality change is further limited by assigning modality without considering biases in PCR amplification, leading to inaccurate modality assignment, such as false bimodal classification (25). Lastly, BRIE2 differential splicing analysis is recommended for comparing PSI values across homogenous, but not, heterogenous cell populations (18). This is because, for a given cell type, BRIE2 imputes alternative splicing events for missing PSI values, using the mean PSI values across the cell population. Moreover, imputed PSI values may not represent actual biological phenomenon (19) and underappreciate the cell-to-cell heterogeneity present in the cell population. Therefore, we sought to apply alternative approaches to compare PSI values between two cell populations and distinguish splicing distributions with similar average PSI values but different PSI distributions, such as bimodal, middle, and multimodal distributions.

To this end, we applied nonparametric tests, namely Kolmogorov-Smirnov (KS), Anderson-Darling (AD), D Test Statistics (DTS) (41), and Wilcoxon rank-sum test, to identify the number of differential splicing events during myoblast differentiation cultured and sequenced at 0- and 72-hours (15). These tests were implemented into MARVEL and identified 39 (KS), 69 (AD), 175 (DTS), and 65 (Wilcoxon rank-sum test) differentially spliced events. Using a comparable cut-off, DTS detected a much higher number of differentially spliced events than other tests, suggesting higher detection sensitivity. However, we observed that differentially spliced events detected by DTS were driven by outlier cells with extremely large or small PSI values relative to most of the cell population (Supplementary Figures 7A-D).

To mitigate differentially spliced events driven by these outlier cells, we applied MARVEL’s bimodal-adjusted modality assignment to identify the events that demonstrated only included to included or excluded to excluded modalities. We set a heuristic threshold of at least 10 cells with PSI values > 0 or < 100 in at least one of two of the cell populations for the differentially spliced events to be retained for downstream analysis (Supplementary Figures 7E-H). We implemented this outlier removal technique into MARVEL. MARVEL’s method removed outliers and retained a higher number of differentially spliced events detected by DTS than other nonparametric tests and BRIE2, which uses the Bayesian model selection method (18) (Supplementary Figure 7I). Because AD and DTS captured most differential splicing events (Supplementary Figure 7J), we combined AD and DTS tests followed by the outlier removal as the default method for MARVEL’s differential splicing analysis. Using a comparable cut-off, MARVEL identified 114 differentially spliced events compared to 73 by BRIE2 (Figure 2J). Next, we investigated whether differentially spliced events detected by MARVEL were biologically relevant to myoblast differentiation. As expected, muscle-related genes were differentially spliced (Supplementary Figures 7K-N). Gene ontology analysis of all differentially spliced genes showed the enrichment of muscle-related pathways when immature myoblasts were differentiated into mature cells (Figure 2K). Moreover, gene ontology analysis of differentially spliced genes detected exclusively by MARVEL additionally identified pathways related to protein translation and localization, and cell cycle pathways (Supplementary Figure 7O).

To assess the generalizability of our method, we performed differentially splicing analysis on single-cell neurons derived from mice induced with multiple sclerosis compared to healthy mice (28). We compared differentially spliced events detected by MARVEL against BRIE2 (in mode 2-diff), which effectively identified differential alternative splicing events for this dataset (18). Using a comparable cut-off, MARVEL identified 248 differentially spliced events compared to 238 by BRIE2 (Figure 2L). Both MARVEL and BRIE2 identified a splicing event that was previously validated using qPCR (28) (Supplementary Figure 7P). Gene ontology analysis of differentially spliced genes detected by MARVEL identified RNA splicing and neuron-related pathways to be enriched, as expected when mice were induced to manifest an autoimmune disease that attacks the nervous system (Figure 2M). Moreover, gene ontology analysis of differentially spliced genes detected exclusively by MARVEL additionally identified lysosome and neurotransmission pathways (Supplementary Figure 7Q).

Taken together, MARVEL identified biologically relevant pathways during muscle cell maturation and in a mouse model with multiple sclerosis. MARVEL complements existing single-cell alternative splicing tools by detecting additional differentially spliced genes and biological pathways.

### MARVEL application for analyzing a plate-based scRNA-seq dataset

After benchmarking MARVEL, we showed the functional features provided by MARVEL to characterize the single-cell alternative splicing landscape in induced pluripotent stem cells (iPSCs) and iPSC-derived endoderm cells (26).

MARVEL detected 13,125 and 5,308 splicing events in iPSCs and endoderm cells. The most prevalent splicing event in iPSCs and endoderm cells was SE, followed by RI, AFE, A3SS, A5SS, ALE, and MXE (Supplementary Figures 8A-B). We investigated whether alternative splicing represented an underappreciated layer of complexity underlying gene expression profile by performing linear dimension reduction analysis using gene expression and PSI values of alternative splicing events. Differentially expressed genes and spliced events robustly distinguished the cell types (Figures 3A and B), whereas non-differentially expressed genes could not separate the cell types (Figure 3C). Interestingly, differential splicing events from non-differentially expressed genes could clearly delineate the two cell types (Figure 3D; Supplementary Figures 8C-I), except for MXE, which was due to the low number of MXE events detected in this analysis (Supplementary Figure 8D). Lastly, all splicing events expressed in both cell types, regardless of whether the events were differentially spliced or not, were also able to separate the cell types (Supplementary Figures 8J).

**Figure 3.**
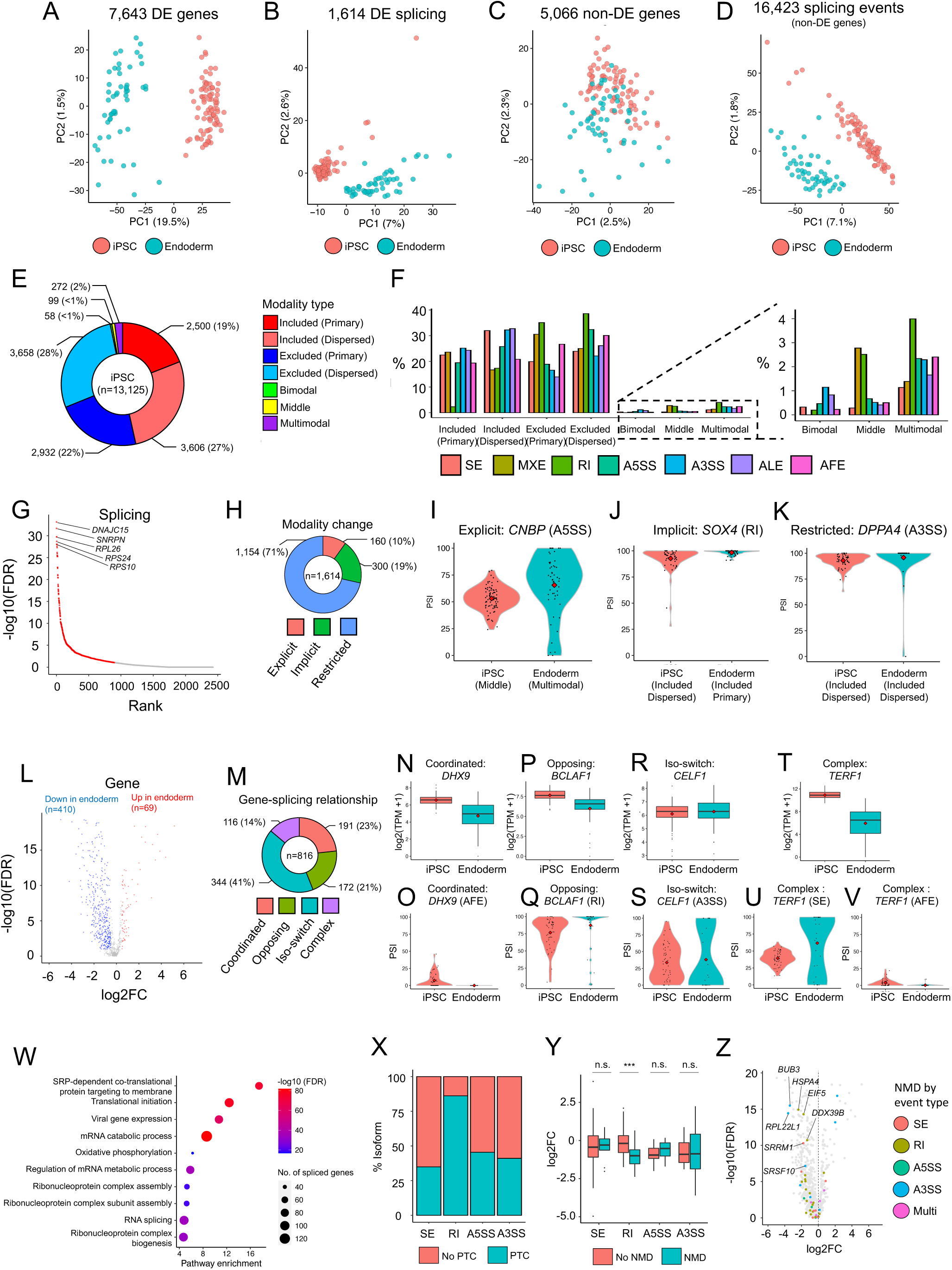
Application of MARVEL on plate-based scRNA-seq data from iPSCs differentiated to endoderm cells. **(A-D)** PCA plots using **(A)** differentially expressed genes, **(B)** differential alternative splicing events, **(C)** non-differentially expressed genes, and **(D)** all alternative splicing events from non-differentially expressed genes. **(E)** The proportion of each modality class in iPSCs **(F)** The proportion of each modality class by splicing event type in iPSCs. **(G)** Ranked list of differentially alternative splicing events identified using MARVEL, comparing PSI distributions between iPSCs and endoderm cells. **(H)** The proportion of each modality dynamic class, i.e., the type of changes in modality of alternative splicing events from iPSCs to endoderm cells. **(I- K)** Representative alternative splicing events of each modality dynamic class **(I)** explicit, **(J)** implicit, and **(K)** restricted. **(L)** Differential expression of genes that were differentially spliced. Blue denotes down-regulated genes (log2FC<-0.5 and FDR<0.10), red denotes up-regulated genes (log2FC>0.5 and FDR<0.10), and grey denotes non-differentially expressed genes (−0.5<log2FC<0.5 or FDR>=0.10). **(M)** The proportion of each gene-splicing relationship class, i.e., the type of changes in average gene expression value relative to change in average PSI value for the corresponding alternative splicing event from iPSCs to endoderm cells. **(N-V)** A representative gene and corresponding alternative splicing event for each gene- splicing relationship class, **(N-O)** coordinated, **(P-Q)** opposing, **(R-S)** isoform switching, and **(T-V)** complex. **(W)** Enrichment scores, FDR values, and gene set sizes of selected biological pathways enriched among differentially spliced genes. **(X)** The proportion of isoforms with PTC introduced by SE, RI, A5SS, and A3SS. **(Y)** Boxplots showing the comparison of log2FC between genes that were not predicted or predicted to be subjected to alternative splicing-mediated NMD. **(Z)** A volcano plot showing differential gene expression analysis between iPSCs and endoderm cells annotated with genes predicted to be subjected to alternative splicing-mediated NMD. FC: fold change; NMD: nonsense-mediated decay; PTC: premature stop codon. *** FDR<0.01 ** FDR<0.05 * FDR<0.1.

Similar to previous studies (19, 26), we next explored PSI distributions (modalities) of individual alternative splicing events identified in iPSCs and endoderm cells. Modality assignment can inform whether a predominant isoform (included and excluded) or both isoforms (bimodal, middle, and multimodal) are expressed and contribute to cellular identity in a cell population. MARVEL assigned distinct modalities to alternative splicing events for each cell type (Figure 3E and Supplementary Figure 8K). Both cell types showed a high proportion of included and excluded modalities (∼97% of all modality types), whereas other modalities (bimodal, middle, and multimodal) showed only ∼ 3%. This is consistent with previous empirical and simulated studies (42, 52). In addition to the original modalities previously proposed (19, 26), MARVEL could stratify the included and excluded modalities into primary and dispersed. In iPSCs (Figure 3E), we observed primary and dispersed modalities constituted 40.9% and 59.1% of the included modality, whereas primary and dispersed modalities constituted 44.5% and 55.5% of the excluded modality. Similar proportions were observed in endoderm cells (Supplementary Figure 8K). Further stratification of modality types by alternative splicing event types showed different proportions of modality types by alternative splicing events (Figure 3F and Supplementary Figure 8L). For example, excluded modality was most prevalent in RI events, constituting 73.6% and 75.2% of all modality types in this splicing event type in iPSCs and endoderm cells. This is consistent with the reported role of intron retention in gene regulation (53).

We performed differential splicing analysis to detect differentially spliced events to understand the splicing dynamics when iPSCs were differentiated into endoderm cells. MARVEL identified 1,614 differential alternative splicing events comprising 816 genes (Figure 3G; Supplementary Tables 2 and 3). Top differentially spliced genes included *DNAJC15*, *SNRPN*, *RPL26*, *RPS24*, and *RPS10*. *DNAJC15* regulates cellular metabolism (54), whereas *SNRPN*, *RPL26*, *RPS24*, and *RPS10* are ribonuclear proteins involved in transcription and translation (55). We selected representative differential alternative splicing events from SE, MXE, RI, A5SS, and A3SS for visual validation using VALERIE (56) (Supplementary Figures 9A-E) (56).

MARVEL further categorized differential alternative splicing events based on modality changes between iPSCs and endoderm cells. We defined modality changes as explicit, implicit, and restricted. Explicit is defined as clear changes within the five main modalities (included, excluded, bimodal, middle, and multimodal). Implicit is defined as changes involving the sub-modalities primary and dispersed. Restricted is defined as no modality changes across the two cell populations. During iPSCs to endoderm cell differentiation, we observed 160, 300, and 1,154 explicit, implicit, and restricted modality changes, respectively, among the differential alternative splicing events (Figure 3H). Notably, most differential alternative splicing events, 1,454 (90%) events, would have been missed if differences in splicing patterns were detected based on explicit modality change alone as used in the previous study (19). Examples of genes that underwent explicit, implicit, and restricted modality change from iPSCs and endoderm cells were *CNBP*, *SOX4*, and *DPPA4* (Figures 3I-K). CNBP is dysregulated in iPSC derived from patients with myotonic dystrophy (57), whereas SOX4 and DPPA4 are transcription factors shown to be dynamically regulated during endoderm induction from iPSCs (58).

General scRNA-seq analysis pipelines perform either differential gene expression or alternative splicing analysis alone but do not integrate both analyses into a single framework. MARVEL allows the integration of differential alternative splicing and gene expression analysis. This enabled us to investigate the changes in alternative splicing relative to changes in gene expression when iPSCs were differentiated into endoderm cells. We identified 816 differentially spliced genes among the 1,614 differentially splicing events. of which, 479 (58%) genes were concurrently differentially expressed (Figure 3L). To explore the relationship between differential genes and alternative splicing changes, MARVEL categorized the changes in gene expression relative to changes in PSI values into coordinated, opposing, isoform switching, and complex (Figure 3M). Coordinated and opposing relationships are defined as changes in gene expression between two cell populations in the same or opposite direction to the change in average PSI values (Figures 3N-Q). For example, DHX9, involved in chromatin remodeling during stem cell differentiation (59), showed coordinated gene-splicing changes, whereby the gene expression and PSI values were decreased from iPSCs to endoderm cells. On the other hand, *BCLAF1* encodes for an anti-apoptotic protein that promotes maintenance and self-renewal of stem cells (60), showed opposing gene-splicing changes, whereby there was a decrease in gene expression from iPSCs to endoderm cells, but PSI values of an RI event were increased. Isoform switching is defined as genes showing differential splicing but not differentially expressed (Figures 3R-S). CELF1, involved in regulating the stability and translation of mRNA during the differentiation process (61), showed no significant difference in gene expression, and mean PSI values in both cell populations were similar, but the overall PSI distribution for an A3SS event was changed from excluded dispersed in iPSCs to bimodal in endoderm cells (explicit modality change). Lastly, a complex relationship involves a combination of coordinated, opposing, and/or isoform switching relationships (Figures 3T-V). For instance, *TERF1*, involved in telomere elongation and maintenance of pluripotency in the iPSCs (62), showed a significant decrease in gene expression in endoderm cells, whereas a SE event of this gene showed a clear modality change from middle to bimodal with a slight increase in PSI values, while a separate AFE splicing event of this gene showed a decrease in PSI values in endoderm cells. Opposing, isoform switching, and complex gene-splicing relationship constituted the majority of the relationships, 632/816 (77%). Therefore, most PSI changes may not be inferred directly from gene expression changes alone. This highlights the value of differential splicing analysis in revealing additional differentially regulated genes.

To assess whether functionally related genes or genes that belong to the same biological pathways are coordinatedly and differentially spliced, we performed gene ontology analysis using 816 differentially spliced genes detected by MARVEL. MARVEL identified 141 significantly enriched pathways among the differentially spliced genes, including pathways related to RNA splicing, gene translation and regulation, and ribonucleoprotein complex formation (Figure 3W). Both RNA splicing and ribonucleoproteins have been shown to regulate stem cell self-renewal and differentiation by modulating protein translation (19, 63).

To understand the functional consequences of alternative splicing of differentially spliced genes, MARVEL predicted nonsense-mediated decay (NMD) for a given alternative splicing event and investigated the relationship between gene expression and alternative splicing-related NMD. We observed RI as alternative splicing events that affected protein-coding transcripts with the highest rate (86%) of introducing premature terminal codons (PTCs), followed by A5SS (46%), A3SS (41%), and SE (35%) (Figure 3X). Only RI-mediated NMD led to a significant decrease in gene expression when iPSCs were differentiated into endoderm cells (Figure 3Y). This is consistent with a previous study that showed a decrease in gene expression by RI-mediated NMD but not NMD mediated by other splicing event types (64). Genes subjected to alternative splicing-related NMD and were concurrently down-regulated in endoderm cells included *BUB3*, *HSPA4*, *EIF5*, *RPL22L1*, *DDX39B, SRRM1*, and the splicing factor *SRSF10* (Figure 3Z). BUB3 is essential for mitotic spindle checkpoint function during cellular proliferation and differentiation (65), whereas HSPA4 represents a class of heat-shock proteins (HSPs) targeting misfolded proteins for degradation (66). EIF5 and RPL22L1 are involved in transcription and translation (67, 68). DDX39B, SRRM1, and SRSF10 regulate RNA splicing (69–71). Taken together, MARVEL links NMD-related splicing changes to gene expression changes to enable prioritization of candidate spliced genes for downstream functional studies (21).

### MARVEL application for analyzing 10x Genomics scRNA-seq dataset

MARVEL also facilitates single-cell alternative splicing analysis for droplet-based scRNA-seq library preparation methods, such as 10x Genomics. To demonstrate the utility of MARVEL, we analyzed scRNA-seq data from the cell differentiation of iPSCs into 10-day-old cardiomyocytes (27). Differential splice junction analysis identified 575 and 243 splice junctions, comprising 539 genes, significantly up or downregulated, in cardiomyocytes relative to iPSCs (Figure 4A; Supplementary Table 4). Differentially spliced genes were enriched in muscle and actin-myosin filament sliding, stem cell differentiation, energy production, and WNT signaling pathways (Figure 4B). Examples of differentially spliced genes included *MYH10*, *ATP5F1C*, and *CBX* (Figures 4C-F) (72). MYH10 is required for proper functioning of the epicardial and formation of coronary vessels (73). *ATP5F1C* encodes a subunit of mitochondrial ATP synthase required for energy production (74). CBX plays a role in chromatin remodeling and neuron development (75).

**Figure 4.**
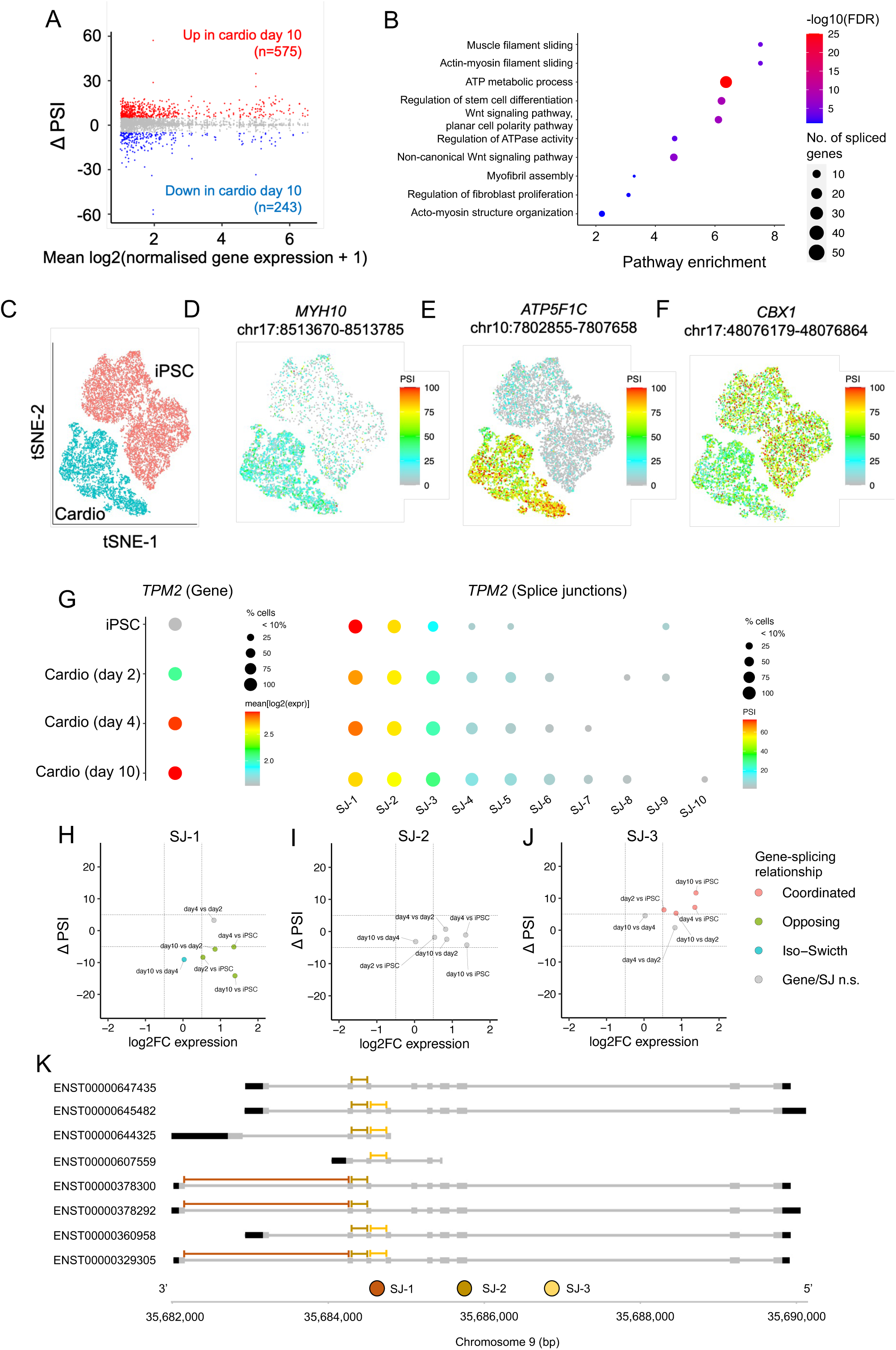
Application of MARVEL on iPSCs differentiated to cardiomyocytes. **(A)** log2FC of splice junction usage in cardiomyocytes relative to iPSCs vs. average splice junction usage across cardiomyocytes and iPSCs. Blue denotes splice junctions down-regulated in cardiomyocytes relative to iPSCs (ΔPSI<-5 and *p*<0.05). Red denotes splice junctions up-regulated in cardiomyocytes relative to iPSCs (ΔPSI>5 and *p*<0.05). Grey denotes splice junctions not differentially spliced between the two cell populations (−5<ΔPSI<5 or *p*>=0.05). **(B)** Enrichment scores, FDR values, and gene set sizes of pathways enriched among differentially spliced genes. **(C-F)** tSNE embeddings generated using 1,160 highly variable genes and consisting of 11,244 iPSCs and 5,937 day-10 cardiomyocytes annotated with **(C)** cell types and muscle-related genes, **(D)** *MYH10*, **(E)** *ATP5F1C*, and **(F)** *CBX1*. **(G)** *TPM2* gene expression and splice junction usage across all cardiomyocyte developmental stages. Splice junctions were arranged in decreasing expression from left to right. **(H-J)** Changes in splice junction usage relative to changes in gene expression levels when comparing more mature cardiomyocytes to less mature cardiomyocytes and iPSCs for the top three highly expressed splice junctions. **(K)** The gene browser shows the relative position of the top three highly expressed splice junctions on *TPM2* protein-coding transcripts.

Next, we investigated gene expression changes relative to splice junction usage changes. Of the 539 genes that were differentially spliced, 19% and 15% of these genes exhibited changes in the same (coordinated) or opposite (opposing) direction relative to splice junction usage, respectively (Supplementary Figure 10A). Genes with a coordinated or opposing relationship with splice junction usage were *VIM* and *UQCRH* (Supplementary Figures 10B-E). *VIM* plays an essential role in maintaining muscle cytoarchitecture and is a reliable marker for muscle cell regeneration (76) while *UQCRH* participates in cardiac muscle contraction (77). More than half (59%) of differentially spliced genes exhibited isoform switching, i.e., differential splice junction usage in the absence of differential gene expression changes, such as the *RBM39* gene that was differentially spliced but not differentially expressed (Supplementary Figures 10F-H). RBM39 is an RNA-binding protein involved in alternative splicing and genetic mutations in *RBM39* are associated with muscle myopathies (78). Last, 7% of genes exhibited a complex relationship with its splice junction usage. Examples of genes with a complex relationship with splice junction usage are *TPM1* and *TPM2* (Supplementary Figures 10I-N). *TPM1* and *TPM2* gene expression were up-regulated in cardiomyocytes relative to iPSCs (Supplementary Figures 10I and L). While one of the splice junctions in both genes exhibited higher expression in cardiomyocytes (Supplementary Figures 10J and M), the other splice junctions exhibited higher expression in iPSCs (Supplementary Figures 10K and N). *TPM1* and *TPM2* are members of the tropomyosin family of highly conserved actin-binding proteins involved in striated and smooth muscle contraction. Genetic mutations in *TPM1* are associated with cardiac hypertrophy, while genetic mutations in *TPM2* were previously reported in patients with congenital myopathy (79, 80).

Most differentially spliced genes, 438 (82%), occurred in the absence (iso-switch) or opposite to splice junction usage changes (opposing relationship) or have a complex relationship with splice junction usage changes. Therefore, most splice junction usage changes cannot be inferred directly from gene expression changes alone.

To further illustrate the intricate relationship between splicing and gene expression profile, we characterized the overall splice junction usage of *TPM2* relative to its corresponding gene expression changes across the developmental stages of cardiomyocytes. We chose *TPM2* for demonstration because its splice junction usage showed a complex relationship relative to its gene expression changes when comparing day-10 cardiomyocytes to iPSCs (Supplementary Figures 10M and N). *TPM2* expression increased from iPSCs to more mature cardiomyocytes (Figure 4G). On the other hand, splice junction-1 (SJ-1; chr9:35682164-35684245) decreased from iPSCs relative to mature cardiomyocytes (Figure 4H), while SJ-2 (chr9:35684316- 35684487) usage was relatively consistent across all developmental stages (Figure 4I). Similar to gene expression changes, SJ-3 (chr9:35684551-35684731) usage increased from iPSCs to mature cardiomyocytes (Figure 4J). The overall splice junction usage across developmental stages decreased from SJ-1 to SJ-2 and SJ-3. We hypothesized that this is due to the 3’-bias inherent in scRNA-seq datasets generated from 3’-bias library preparation methods. To this end, we implemented a gene browser visualization function in MARVEL to inspect the specific location of splice junctions of interest relative to the transcripts. Indeed, SJ-1 was located on the most 3’-end of the transcripts, followed by SJ-2 and SJ-3 (Figure 4K). Therefore, the end-bias inherent in single-cell library preparation methods would be taken into account during single-cell alternative splicing analysis.

## DISCUSSION

We have developed MARVEL to address key issues in single-cell alternative splicing analysis and enable transcriptome-wide characterization of the alternative splicing dynamics in scRNA-seq datasets. We benchmarked MARVEL against the existing alternative splicing analysis tools and demonstrated the utility of MARVEL using for datasets generated from the plate- and droplet-based methods. We summarized the available features of MARVEL compared to other single-cell alternative splicing analysis tools shown in Supplementary Table 1.

MARVEL employed a splice junction-based approach to estimate the PSI directly from splice junction reads that reflect true biological phenomena (19). PSI values estimated by using probabilistic frameworks, such as BRIE, have shown bias in PSI estimation and underestimated cell-to-cell heterogeneity at low coverage (17,81,82). We showed that MARVEL had a better cell-to-cell and cell-to-bulk correlation of PSI values in homogenous cell lines than other approaches.

Most single-cell alternative splicing analysis tools constrained PSI quantification to only an exon-skipping splicing event. Nevertheless, other splicing event types also contribute to the cellular phenotype. For example, aberrant intron retention in cancer leads to abnormal proteins presented on the tumor surface as neoantigens, which may be amenable to immunotherapy (22). Alternative 3’ splice sites are preferentially mis-spliced by mutant splicing factor *SF3B1* (47, 83). Therefore, MARVEL has been developed to include PSI quantification for all main exon-level splicing event types, comprising SE, MXE, RI, A5SS, A3SS, AFE, and ALE. MARVEL also requires less processing time and lower memory usage than other tools.

PSI values reflect the percentage of splice junction reads supporting the alternative exons and are therefore represented by any values between 0-100. Song *et al.* previously introduced the concept of “modality” to categorize the PSI distribution for a given alternative splicing event into discrete categories (19). The classes of modalities were included, excluded, bimodal, middle, and multimodal. MARVEL introduces primary and dispersed sub-modalities for included and excluded modalities. We showed that ∼50% of included and excluded comprised of primary sub-modality, whereas another ∼50% showed dispersed sub-modality, suggesting that MARVEL could increase the current repertoire of modality classes and provide a finer distinction between the different PSI distributions.

A significant proportion of bimodality may have been misclassified (25). We tabulated a catalog of true and false bimodal alternative splicing events previously validated using qPCR, smFISH, and inferred mRNA counts (15,19,25,26). We identified key features that distinguished true from false bimodality. These features were incorporated into MARVEL to identify and adjust for the false bimodal class, leading to more accurate modality classification and modality change detection between different cell populations.

Current approaches for differential alternative splicing analysis in single cells include a comparison of two cells at a time or detection of modality changes between cell populations. For the former approach, comparing all possible cell pairs is impractical when the number of cells becomes large (17). For the latter approach, changes in splicing patterns are defined on modality changes across different cell populations, such as included to excluded modality change (19). This approach may miss changes in splicing patterns that do not involve any modality change. MARVEL incorporated the statistical framework Anderson-Darling and D Test Statistic (41) combined with the bimodal-adjusted modality assignment to enable unbiased evaluation of the differences in PSI distribution across different cell populations. We showed that 90% of differential alternative splicing events identified by MARVEL, when iPSCs were differentiated into endoderm cells, demonstrated no explicit change in PSI modality. These events would have been missed based on the original modalities proposed by Song *et al.* (19).

Current single-cell analysis tools offer only gene or alternative splicing analysis exclusively (17,19,84). Nevertheless, alternative splicing and gene expression changes may be intricately linked. MARVEL integrates alternative splicing and gene expression to study the relationship between alternative splicing and gene expression changes across different cell populations. For example, comparative analysis between iPSCs and endoderm cells demonstrated that only about 23% of differentially spliced genes showed gene expression changes that occurred in the same direction as the corresponding PSI changes. The remaining differentially spliced genes occurred in the opposite or the absence of gene expression changes. This reaffirms the complex relationship underlying gene expression and alternative splicing.

Current single-cell alternative splicing analysis tools fall short in providing context to understand the functional consequence of alternative splicing. Alternative splicing represents one of many mechanisms by which gene expression is regulated. We incorporated nonsense-mediated decay prediction (NMD) as a functional annotation feature, in addition to gene ontology analysis, into MARVEL. MARVEL can predict whether the insertion of a given alternative exon subjects the corresponding isoforms to NMD or not. It compares gene expression levels between genes that are subjected to NMD. We showed that increased intron retention decreased gene expression levels when iPSCs were differentiated into endoderm cells. This is reminiscent of the complex, in this case opposing relationship between alternative splicing and gene expression changes. This finding is in line with a previous report demonstrating that only intron retention, but not other splicing event types, was associated with decreased gene expression levels using long-read RNA-sequencing in *SF3B1*-mutated chronic lymphocytic leukemia patients (64).

We extended MARVEL’s framework to enable integrated gene and alternative splicing analysis in the dataset generated from a droplet-based platform. MARVEL was able to identify differential splice junction usage enriched in muscle-, neuron-, and heart-related pathways in iPSCs differentiated to cardiomyocytes. Moreover, only 19% of differentially spliced genes demonstrated the same directional changes in splice junction usage and gene expression. This is consistent with the intricate relationship between alternative splicing changes and gene expression changes revealed by the plate-based analysis. Lastly, MARVEL enables single-cell visualization of splice junction usage on linear or non-linear dimensionality reduction to verify differential splicing junction usage across different cell populations.

Our study provides a comprehensive computational framework to characterize alternative splicing dynamics at single-cell resolution. As far as we are aware, MARVEL is the only single-cell alternative splicing computational tool to enable alternative splicing analysis on scRNA-seq data generated from the plate- and droplet-based library preparation methods. MARVEL supports the integration of gene-level expression and alternative splicing analysis. For alternative splicing analysis, MARVEL provides end-to-end features to characterize single-cell alternative splicing landscape, starting from alternative splicing event validation, percent spliced-in quantification, modality assignment and correction, differential splicing analysis, to the functional annotation using gene ontology and nonsense-mediate decay prediction. We anticipate MARVEL to be prospectively applied to single-cell datasets generated from various settings (e.g., health and disease states) to reveal novel biological insights.

## DATA AVAILABILITY

MARVEL is available on GitHub: https://github.com/wenweixiong/MARVEL. The software tutorial containing the pre-processed data and codes to reproduce the figures related to the application of MARVEL on plate- and droplet-based RNA-sequencing data are available at https://wenweixiong.github.io/MARVEL_Plate.html and https://wenweixiong.github.io/MARVEL_Droplet.html, respectively. All data sources included in this study are publicly available.

## ACKNOWLEDGMENTS

The authors would like to thank the members of the WIMM Centre for Computational Biology for testing the MARVEL package and providing useful input.

---

**Supplementary Figure 1.**
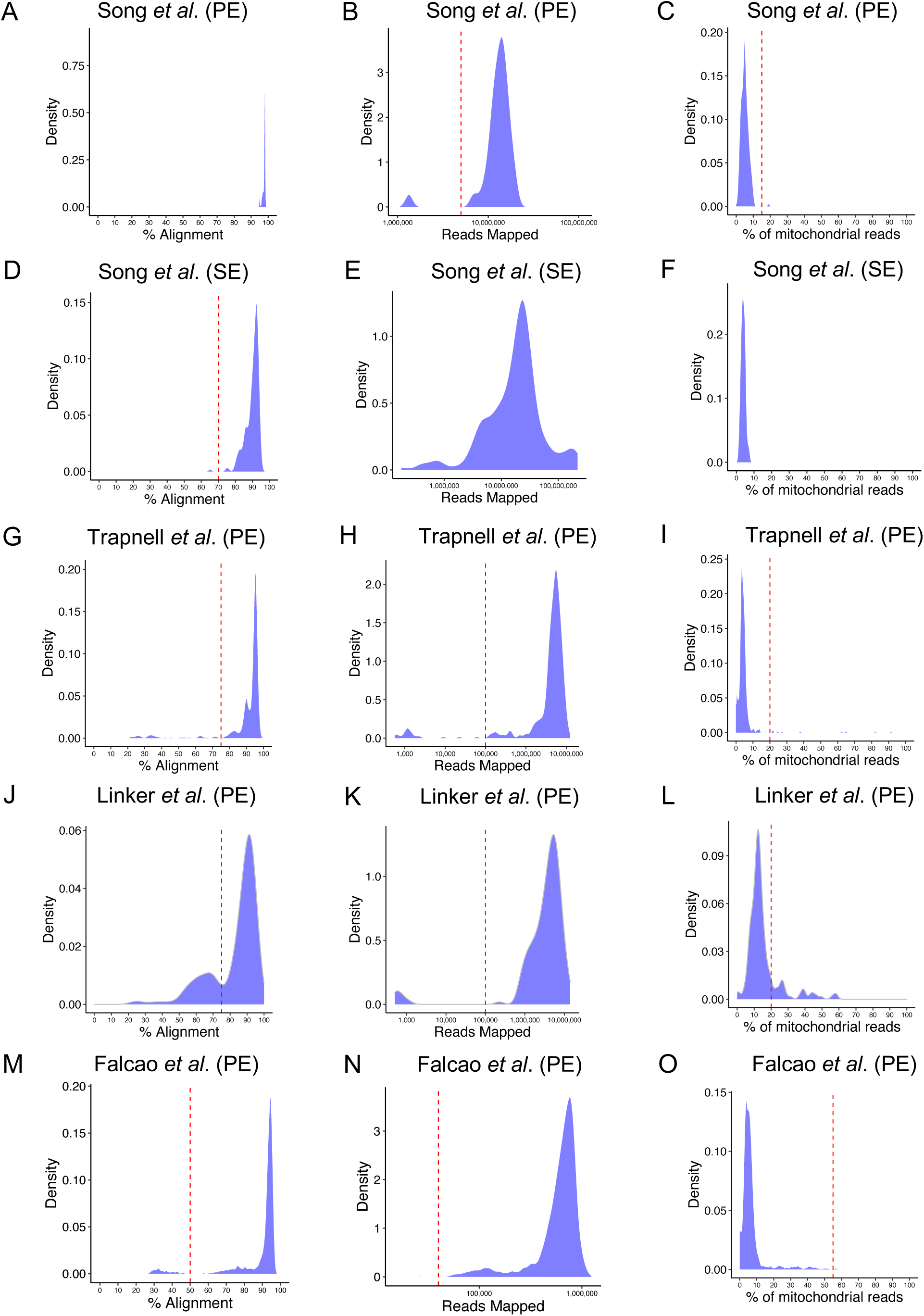
QC to identify high-quality cells for downstream analyses for plate-based scRNA-seq datasets. **(A-L)** Sequencing QC using total reads mapped, alignment rate, and percentage of mitochondrial reads contribution to filter for high-quality cells in **(A-F)** Song *et al.*, **(G-I)** Trapnell *et al.*, **(J-L)** Linker *et al.*, and **(M-O)** Falcao *et al*. dataset. For total reads mapped and alignment rate, cells above the threshold denoted by the red dashed line were considered high-quality. For the percentage of mitochondrial reads contribution, cells below the threshold denoted by the red dashed line were considered high-quality. Only cells meeting all three criteria were included for downstream analyses.

**Supplementary Figure 2.**
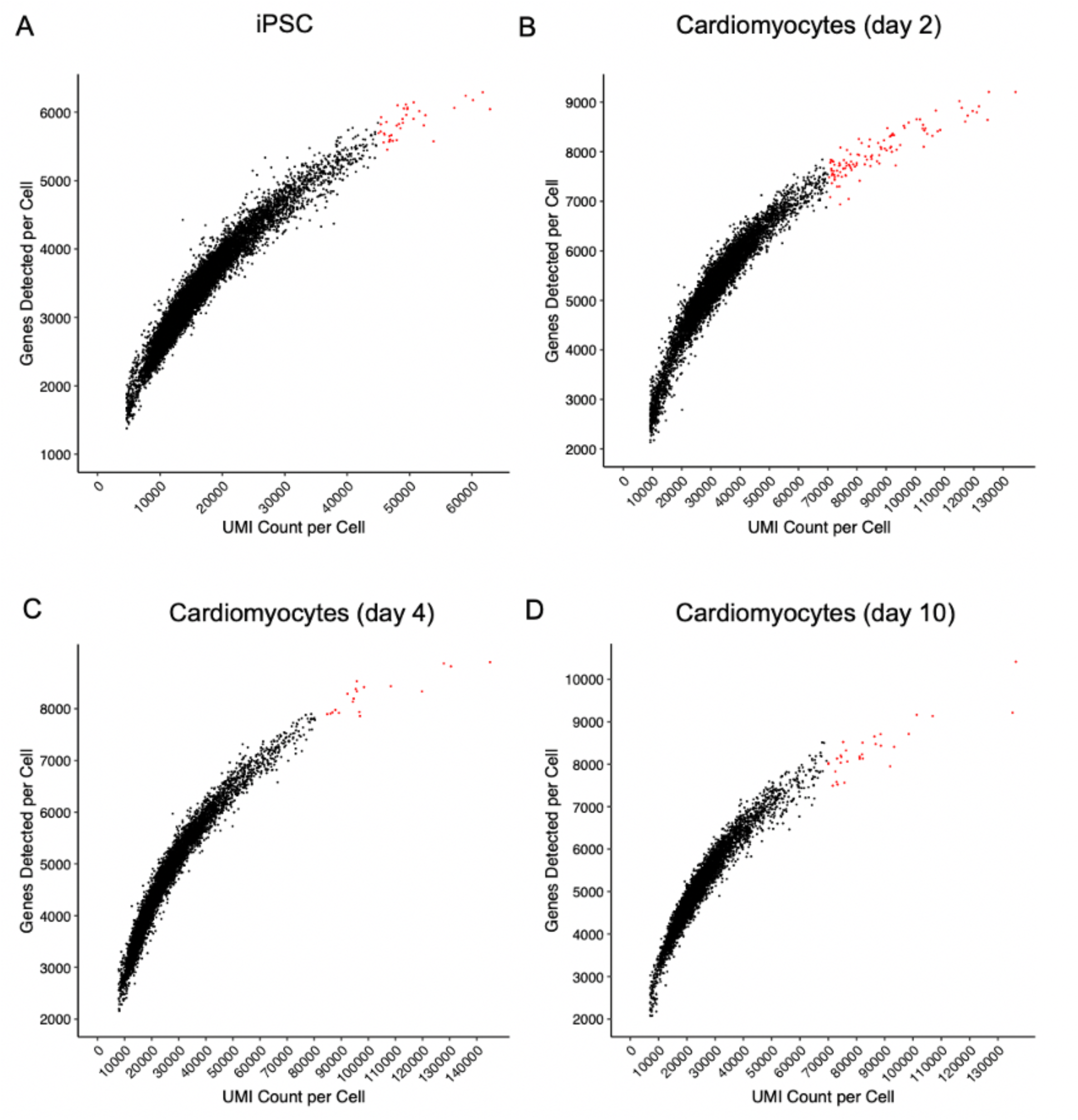
QC to identify high-quality cells for downstream analyses for the droplet-based scRNA-seq dataset. **(A-D)** QC of single cells based on the number of genes detected and number of UMIs for **(A)** iPSCs, and cardiomyocytes at days **(B)** 2, **(C)** 4, and **(D)** 10. The red cells were excluded from downstream analyses based on UMI counts and the number of detected genes. UMI: unique molecular identifier.

**Supplementary Figure 3.**
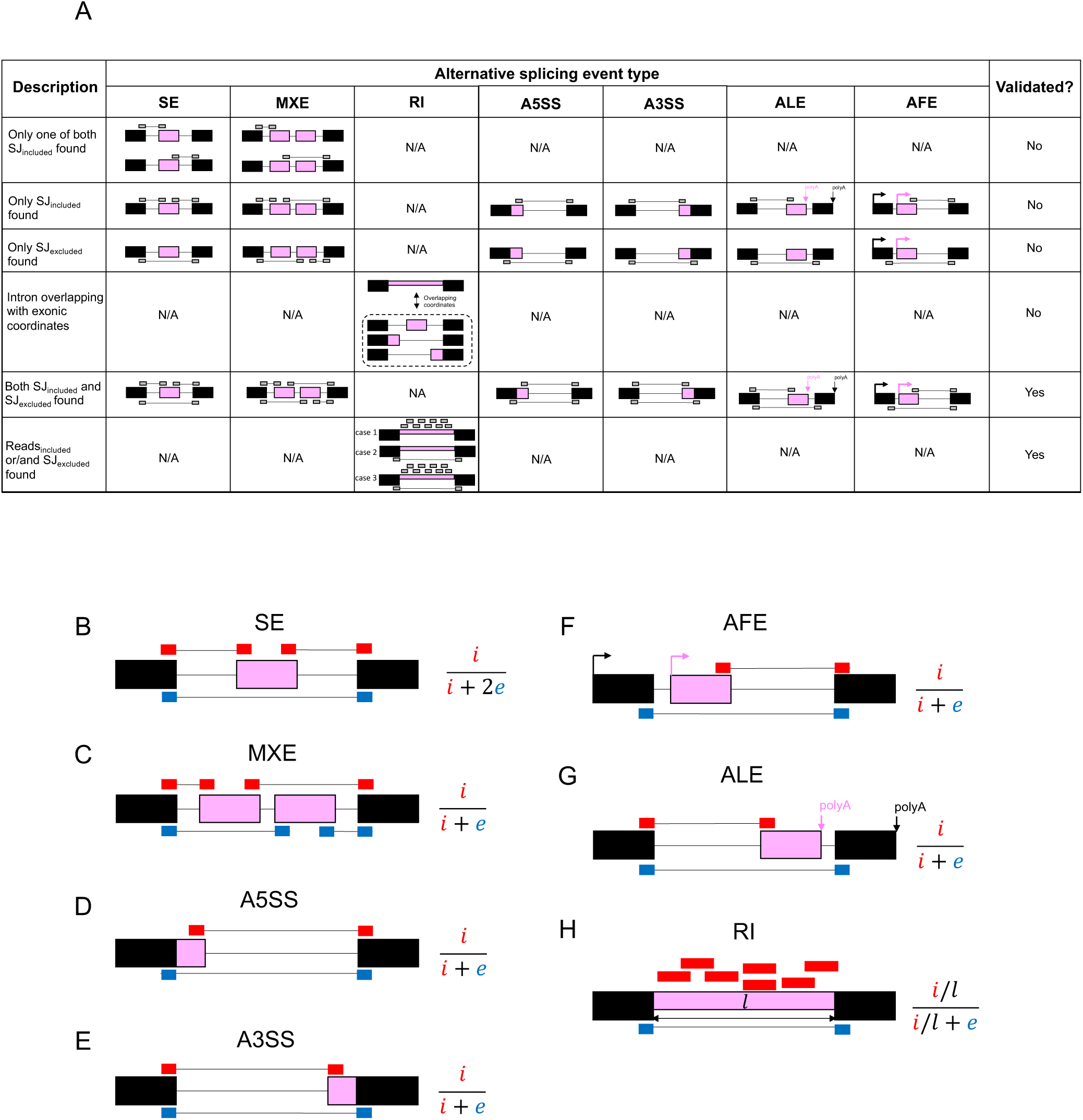
Alternative splicing event validation and PSI formulas. **(A)** Validation of alternative splicing events using splice junction reads to select high-quality alternative splicing events for PSI estimation. **(B-F)** The formulas to estimate the PSI values for **(B)** SE, **(C)** MXE, **(D)** A5SS, **(E)** A3SS, **(F)** AFE, **(G)** ALE, and **(H)** RI. *i* denotes the number of splice junction reads supporting the alternative exon(s) in pink. *e* denotes the number of splice junction reads supporting the constitutive exon(s) in black but skipping the alternative exon(s). *l* denotes the intron length.

**Supplementary Figure 4.**
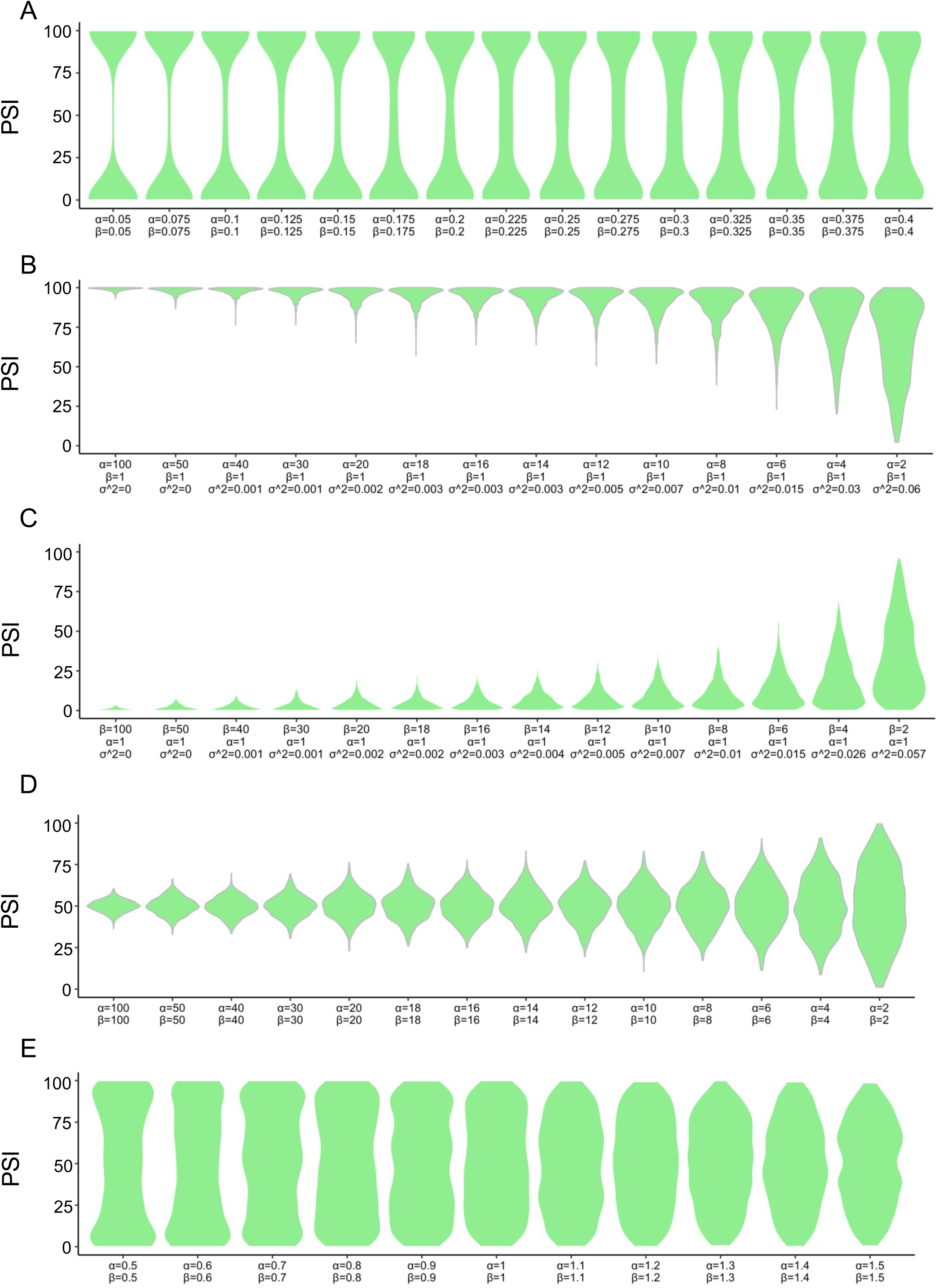
The simulation of PSI values in different PSI distributions for each modality class. **(A-E)** PSI distributions corresponding to **(A)** bimodal, **(B)** included, **(C)** excluded, **(D)** middle, and **(E)** multimodal. The PSI distributions were modeled using the beta distribution, and the corresponding α and β parameters were estimated using the maximum likelihood approach.

**Supplementary Figure 5.**
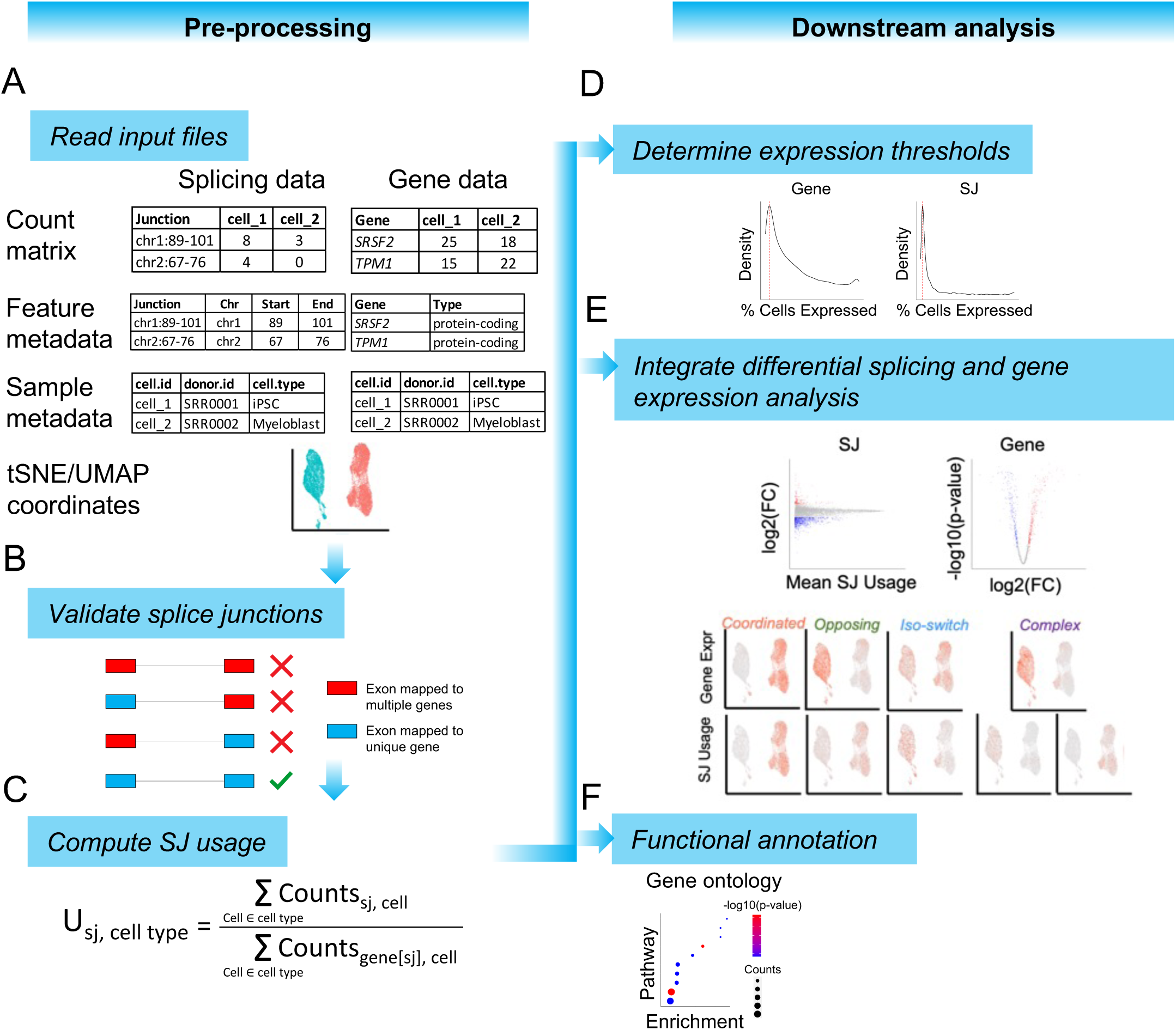
MARVEL workflow for single-cell alternative splicing analysis in a droplet-based scRNA-seq dataset. **(A-C)** Pre-processing of splice junction and gene expression data. **(A)** Inputs for MARVEL include splice junction and gene count matrix, normalized gene expression matrix, and splice junction, gene and sample metadata, and dimension reduction coordinates. **(B)** Only splice junctions, in which both exons are mapped to the same unique gene are retained. **(C)** For a given cell population, the splice junction usage is computed for the high-quality splice junctions identified in **(B)** as the total splice junction counts divided by the total corresponding gene counts. **(D-E)** Downstream analyses using the computed splice junction usage and gene expression values. **(D)** Splice junction and gene expression distributions across a given cell population. Red dotted lines correspond to the percentage of cells in which most splice junctions and genes are expressed and can be used as thresholds for sub-setting splice junctions and genes for differential analysis. **(E)** Integrative differential splice junction and gene expression analysis allows for investigating changes in splice junction usage relative to changes in gene expression across different cell populations. **(F)** Pathway enrichment analysis to identify gene sets that are coordinatedly spliced. iPSC: Induced pluripotent stem cells.

**Supplementary Figure 6.**
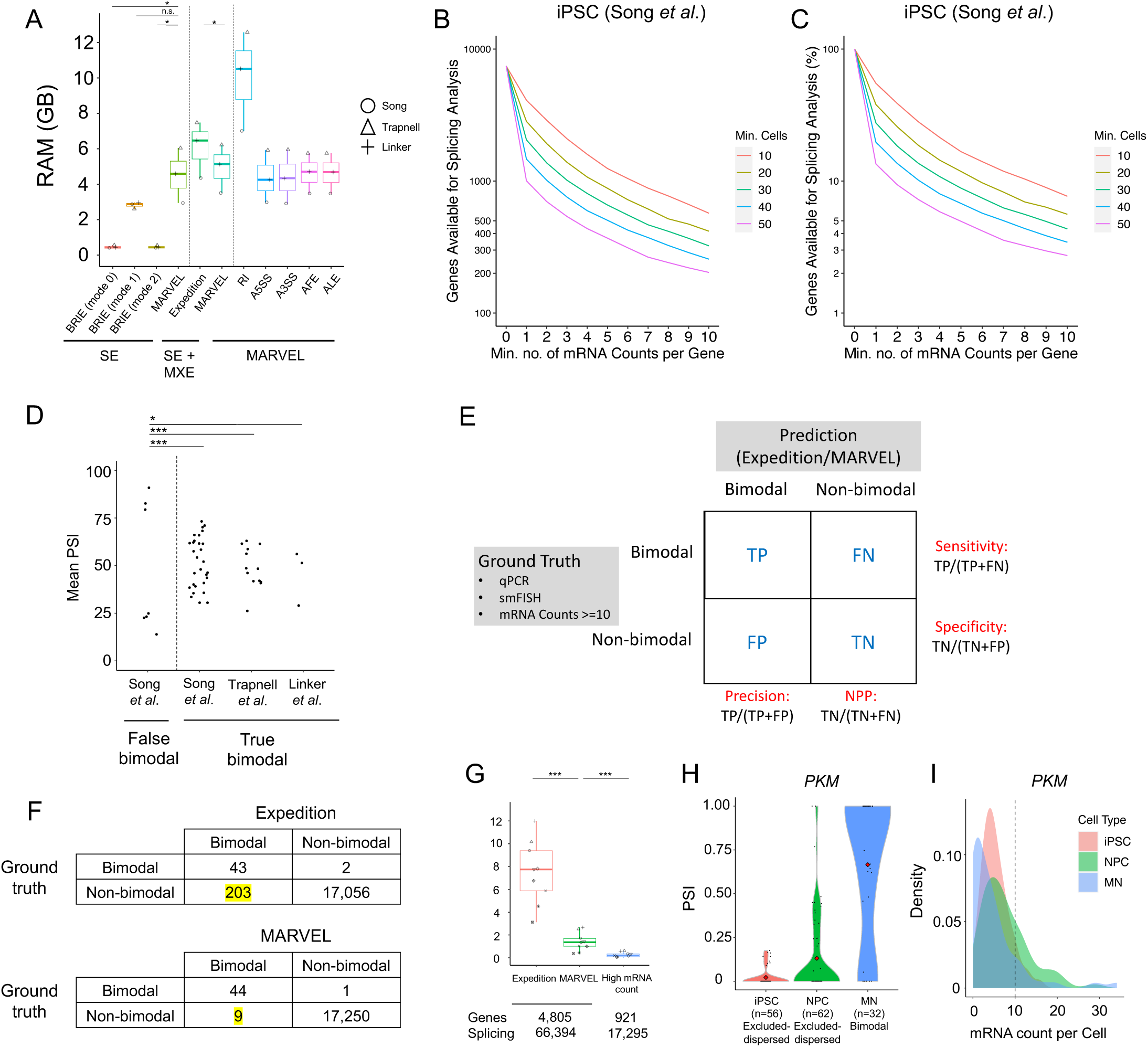
Benchmarking MARVEL against existing computational tools for estimating PSI values and modality assignment. **(A)** The use of RAM in GB to compute PSI values for 1,000 splicing events for each splicing event type in three datasets. **(B-C)** In iPSCs, the **(B)** number or **(C)** percentage of genes for alternative splicing analysis at different mRNA count thresholds and the different minimum number of cells thresholds (sample size). **(D)** The average PSI values of false and true bimodal distributions. **(E-F)** The confusion matrix comparing the number of bimodal and non-bimodal classifications assigned by Expedition and MARVEL compared to that of the ground truth consisting of 17,304 false and true bimodal distributions. **(G)** The percentage of alternative splicing events assigned as bimodal distribution by Expedition, MARVEL, and high mRNA count approach. **(H)** The PSI distribution for *PKM* alternative splicing event in iPSCs, NPCs, and MNs. **(I)** The mRNA count distribution for *PKM* in iPSCs, NPCs, and MNs. RAM: Random Access Memory; GB: Gigabyte. *** FDR < 0.01 ** FDR < 0.05 * FDR < 0.1.

**Supplementary Figure 7.**
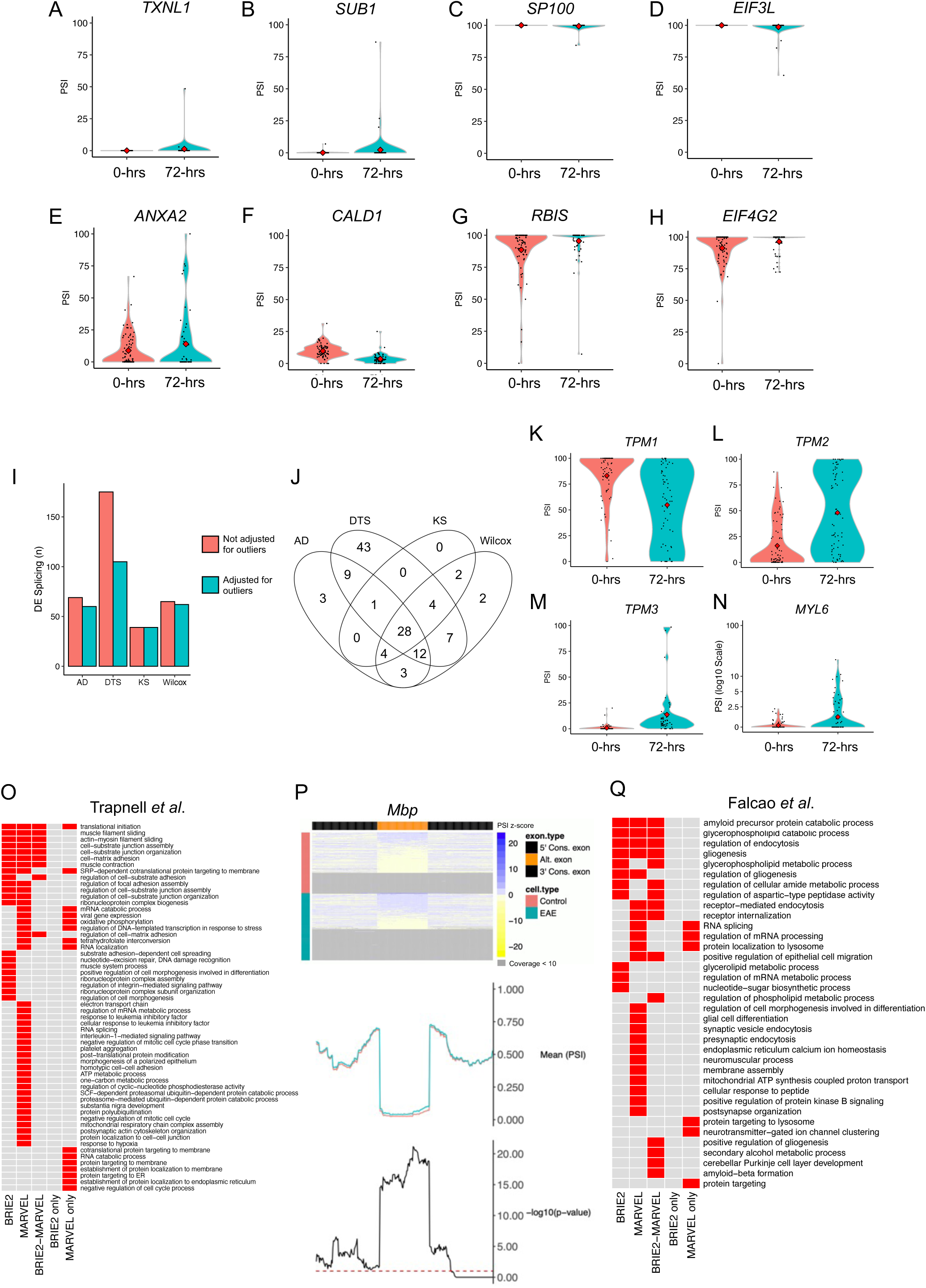
Benchmarking MARVEL against existing computational tools for differential splicing analysis. **(A-D)** Representative examples of splicing events, identified as differentially spliced by DTS, driven by the small number of cells (outliers) with PSI values of **(A-B)** >0 or **(C-D)** <1 when the modality change is from excluded to excluded or included to included, respectively. **(E-H)** Representative examples of splicing events that were considered as differentially spliced when the number of cells with PSI values of **(E-F)** >0 or **(G-H)** <1 when in either cell group is above a user-defined threshold, in this study is 10, when the modality change is from included to included or excluded to excluded, respectively. **(I)** The number of differentially spliced events detected between 72- vs. 0-hrs myoblast before and after removing events driven by outlier cells. **(J)** The number of differentially spliced events detected by AD, DTS, KS, and Wilcoxon rank-sum tests between 72- vs. 0-hrs myoblast. **(K-N)** Representative examples of muscle-related differentially spliced genes detected by MARVEL. **(O)** Identified pathways enriched among differentially spliced genes between 72- vs. 0-hrs myoblast among differentially spliced events identified by “BRIE2” (all spliced genes identified by BRIE2), “MARVEL” (all spliced genes identified by MARVEL), “BRIE2-MARVEL overlap” (spliced genes detected by both BRIE2 and MARVEL), “BRIE2-only” (spliced genes identified by BRIE2 but not MARVEL), and “MARVEL-only” (spliced genes identified by MARVEL but not BRIE2). **(P)** The visual validation of *Mbp* exon 2 shows differentially spliced between EAE and control mice by MARVEL. This exon2 was also previously validated using qPCR by the original study. **(Q)** Identified pathways enriched among differentially spliced genes between EAE and control mice among differentially spliced events identified by “BRIE2” (all spliced genes identified by BRIE2), “MARVEL” (all spliced genes identified by MARVEL), “BRIE2-MARVEL overlap” (spliced genes detected by both BRIE2 and MARVEL), “BRIE2-only” (spliced genes identified by BRIE2 but not MARVEL), and “MARVEL-only” (spliced genes identified by MARVEL but not BRIE2). AD: Anderson-Darling; DTS: D Test Statistic; EAE: experimental autoimmune encephalomyelitis; KS: Kolmogorov-Smirnov; hrs: hours; PSI: percent spliced-in; qPCR: quantitative polymerase chain reaction.

**Supplementary Figure 8.**
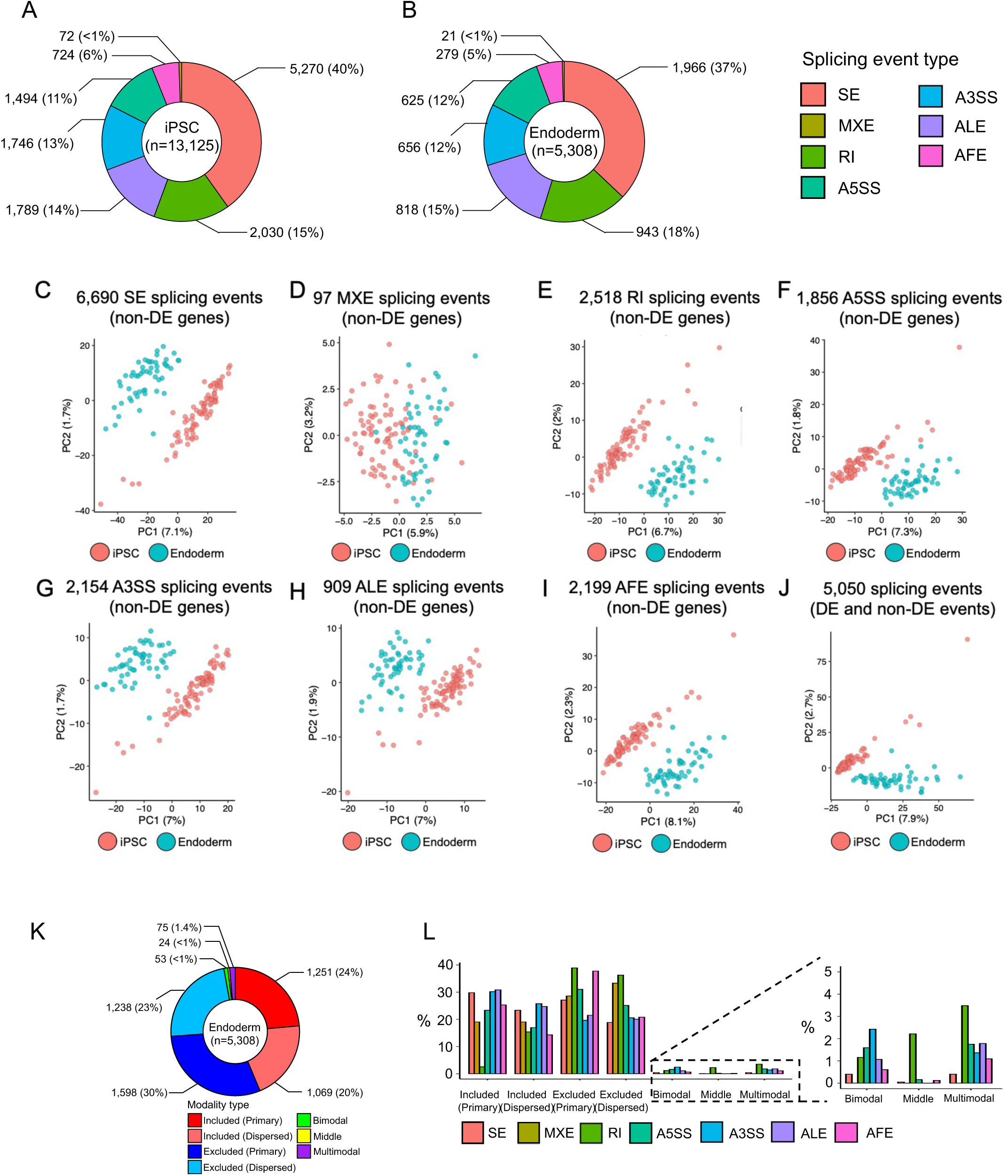
Application of MARVEL on iPSCs differentiated to endoderm cells. **(A-B)** The proportion of expressed alternative splicing event types in **(A)** iPSCs and **(B)** endoderm cells. **(C-J)** Dimension reduction analysis with PCA using **(C)** SE, **(D)** MXE, **(E)** RI, **(F)** A5SS, **(G)** A3SS, **(H)** ALE, and **(I)** AFE splicing events of non-differentially expressed genes, and **(J)** all splicing events regardless of if there were differentially spliced or not. **(K)** The proportion of each modality class in endoderm cells. **(L)** The proportion of each modality class by splicing event type in endoderm cells. A3SS: alternative 3’ splice site; A5SS: alternative 5’ splice site; AFE: alternative first exon; ALE: alternative last exon; MXE: mutually exclusive exons; RI: retained-intron; SE: skipped-exon.

**Supplementary Figure 9:**
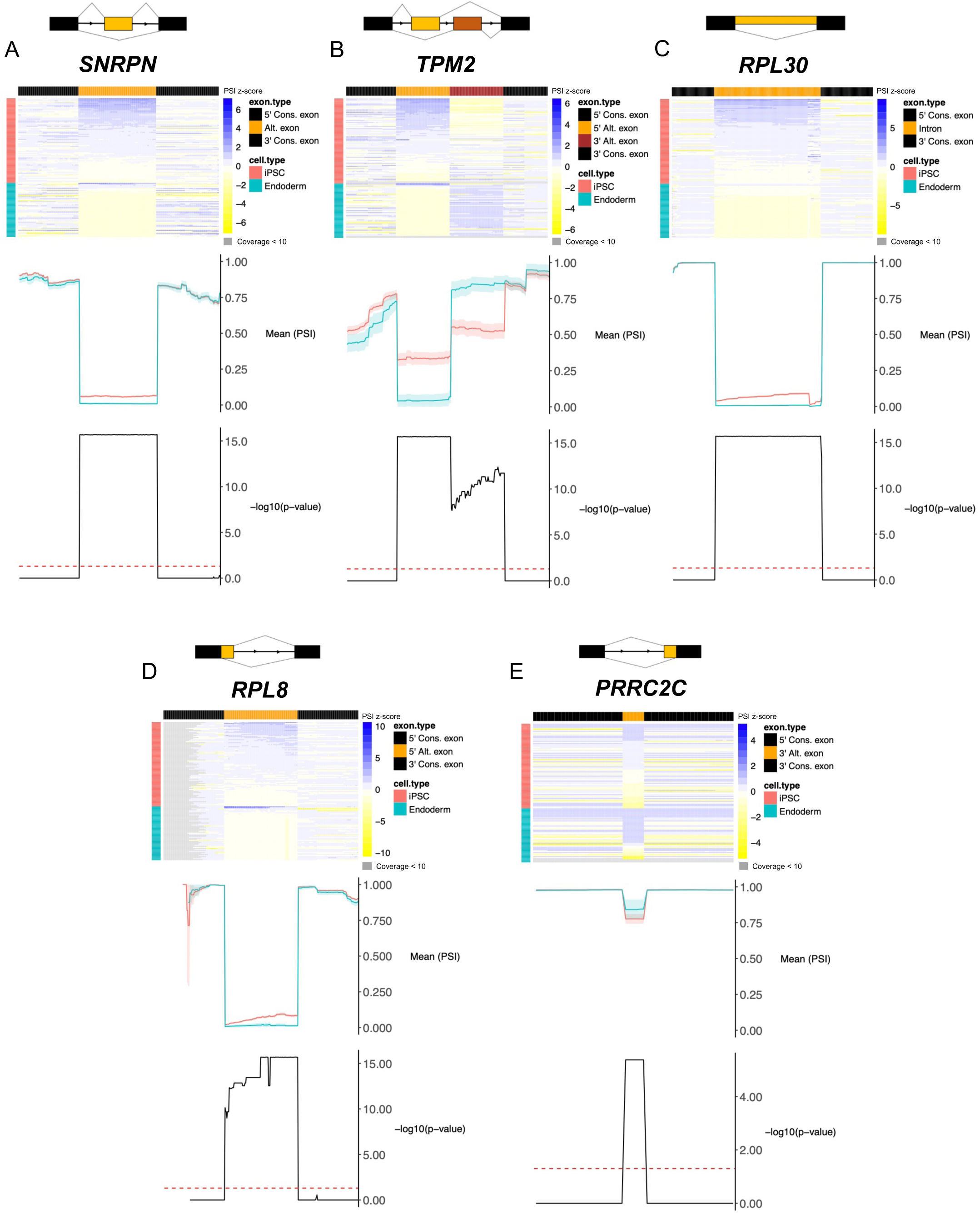
Visual validation of differential alternative splicing events detected by MARVEL using VALERIE. **(A-E)** Representative examples of genes for each splicing event, **(A)** SE, **(B)** MXE, **(C)** RI, **(D)** A5SS, and **(E)** A3SS visually inspected using VALERIE. A3SS: alternative 3’ splice site; A5SS: alternative 5’ splice site; MXE: mutually exclusive exons; RI: retained-intron; SE: skipped-exon.

**Supplementary Figure 10.**
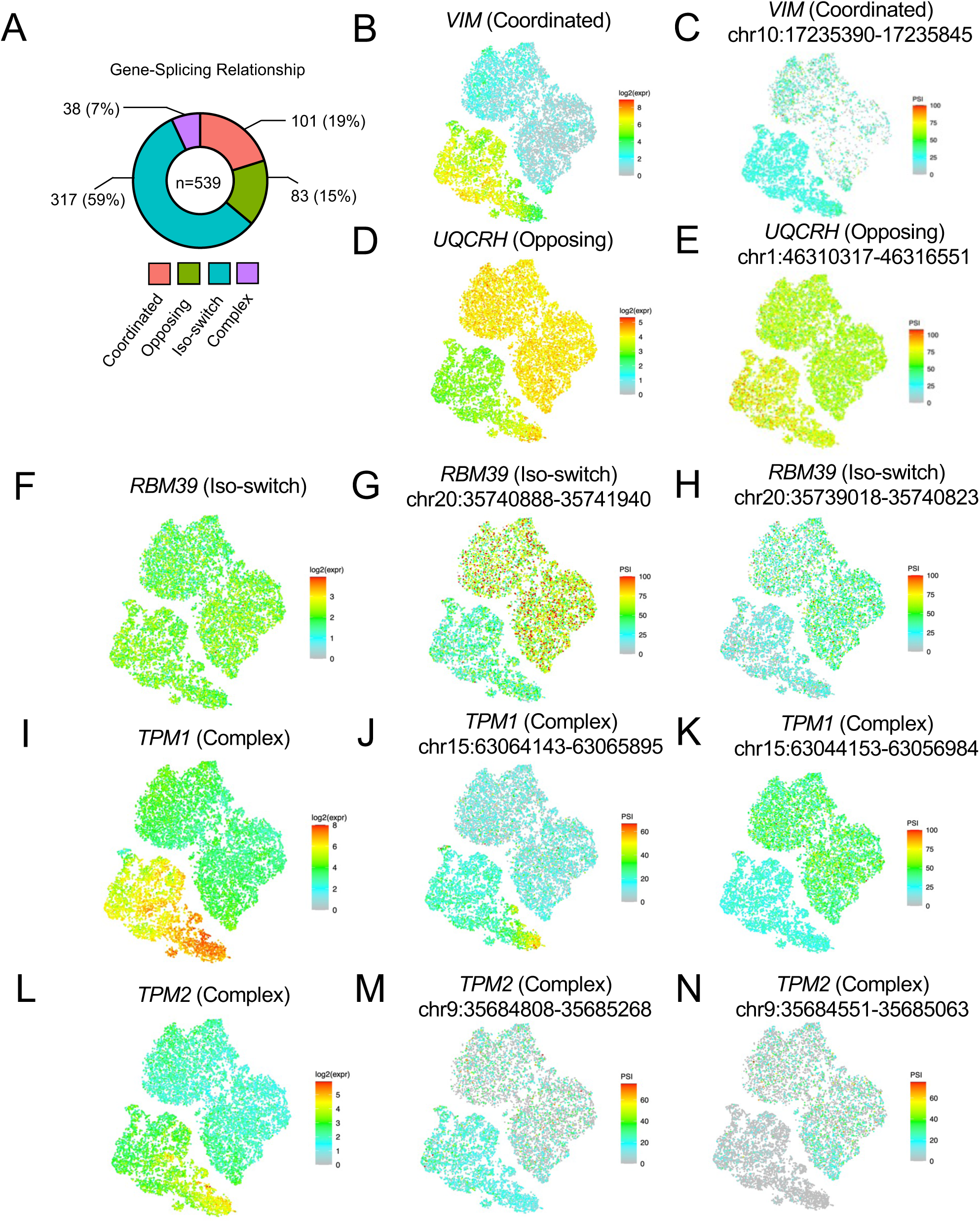
Gene expression and splice junction usage relationship of differentially spliced genes identified from iPSCs differentiated to day-10 cardiomyocytes. **(A)** The proportion of each gene-splicing relationship class, i.e., change in average gene expression value relative to change in average splice junction usage value for the corresponding splice junction from iPSCs differentiated to cardiomyocytes. **(B-K)** Representative gene expression and corresponding splice junctions for each gene-splicing relationship class, **(B-C)** coordinated, **(D-E)** opposing, **(F-H)** isoform switching, and **(I-N)** complex. Gene expression levels are indicated by the log2 scale. The splicing rate is indicated by the PSI scale. The genomic coordinates are also indicated for the respective splice junctions.

