## Supplementary Table 1 for "MARVEL: An integrated alternative splicing analysis platform for single-cell RNA sequencing data"

**Supplementary Table 1. Analysis features provided by MARVEL compared to BRIE2 and Expedition**

| **Feature** | **BRIE2** | **Expedition** | **MARVEL** |
| --- | --- | --- | --- |
| **PSI estimation** |  |  |  |
| SE | / | / | / |
| MXE | x | / | / |
| RI | x | x | / |
| A5SS | x | x | / |
| A3SS | x | x | / |
| AFE | x | x | / |
| ALE | x | x | / |
| **Modality classification** |  |  |  |
| Main modalities | x | / | / |
| Sub-modalities | x | x | / |
| False modality correction | x | x | / |
| **Differential expression analysis** |  |  |  |
| Differential splicing analysis | / | x | / |
| Differential gene expression integration | x | x | / |
| **Functional annotation** |  |  |  |
| Gene ontology | x | x | / |
| Nonsense-mediated decay prediction | x | x | / |
| **RNA-velocity** | / | x | x |
| **RNA-sequencing analysis** |  |  |  |
| Plate-based (e.g., Smart-seq2) | / | / | / |
| Droplet-based (e.g., 10x Genomics) | x | x | / |
